# NMR assignment and functional characterization of biotechnologically modified xanthan

**DOI:** 10.64898/2026.10.28.762522

**Authors:** Julian Wegner, Pia Lanvers, Kristoffer Johansen, Moritz Gansbiller, Bjørn E. Christensen, Jochen Schmid, Gaston Courtade

## Abstract

Xanthan gum is one of the most important industrial microbial polysaccharides, yet comprehensive molecular-level structural characterization remains limited because of its high molecular weight, conformational rigidity, and severe spectral overlap in nuclear magnetic resonance spectroscopy. Here, we present a near-complete ^1^H/^13^C NMR chemical shift assignment of the xanthan repeating unit by combining multidimensional NMR spectroscopy with targeted glycosyltransferase knockout strains and controlled enzymatic depolymerization. Comparative analysis of xanthan from the wild-type strain and the *ΔgumI* and *ΔgumK* variants enabled unambiguous assignment of β-mannose and glucuronic acid resonances and, by extension, the remaining sugars, supporting future structural studies of engineered xanthan variants. In addition, we investigated how side-chain composition affects xanthan properties by analyzing genetically modified variants lacking specific acetyl, pyruvyl, or sugar residues. Temperature-dependent ^1^H NMR determined order-disorder transition temperatures and revealed that removing pyruvate or negatively charged side-chain components increased thermal stability, whereas losing the terminal β-mannose destabilized the ordered conformation. Together, these results establish a robust NMR assignment for xanthan and provide new insights into how individual side-chain substituents influence the structural stability of this polysaccharide, facilitating the rational characterization of engineered xanthan variants and structurally closely related polysaccharides.

**Graphical abstract:** 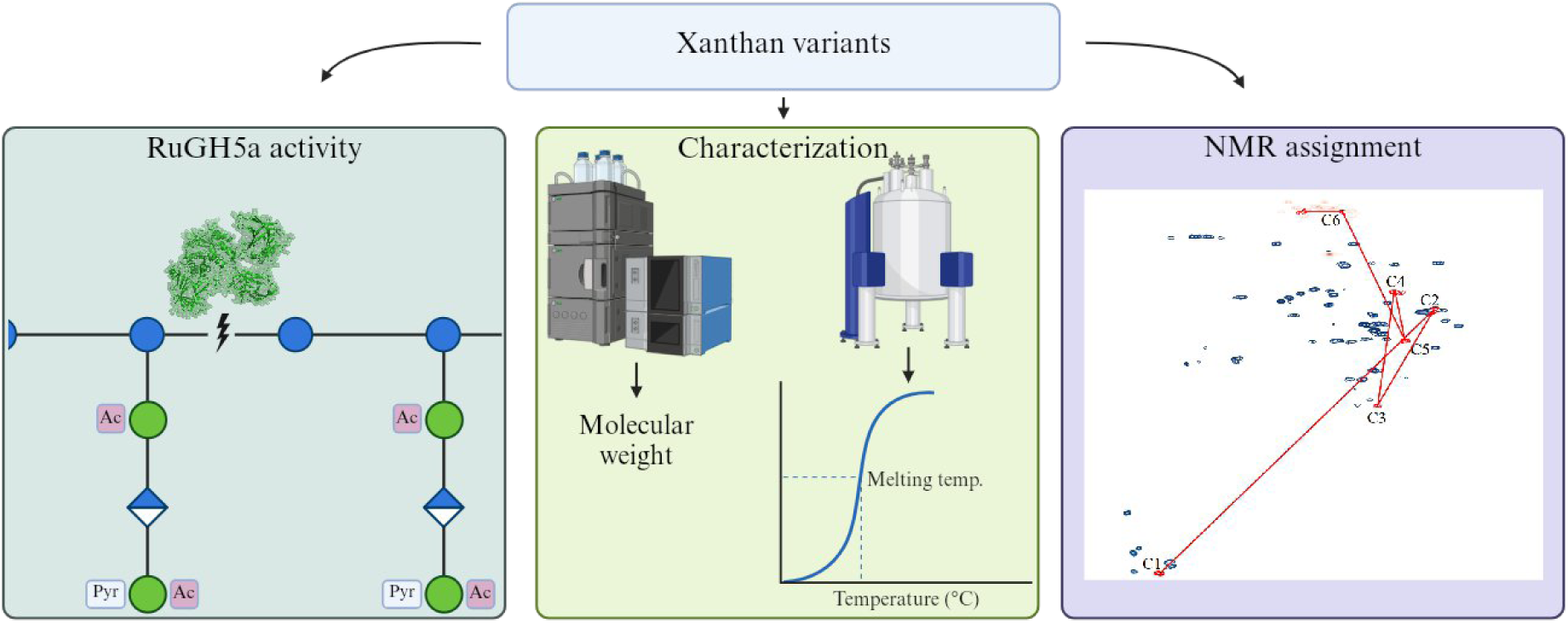

## 1. Introduction

Bacteria produce extracellular polysaccharides (EPS) with an enormous range of chemical architectures. Each EPS is tailored to specific biological roles, from evading host immunity to forming resilient biofilms (Whitfield, 1988). This structural diversity also makes EPS attractive targets for biotechnological modification (Whitfield, 1988).

Among microbial polysaccharides, xanthan gum (xanthan), an EPS produced by *Xanthomonas campestris pv. campestris* (*Xcc*) occupies a dominant position in industrial applications, with annual production exceeding 165,000 tons (Sutherland, 1998; Vorhölter et al., 2008; Wilson et al., 2020). Around two-thirds of the worldwide demand stems from the food sector, with the remaining share used in petroleum-related applications and various technical industries (Petri, 2015; Vorhölter et al., 2008). These widespread applications stem from xanthan’s exceptional viscosity and distinctive rheological characteristics (Rudolph, 1993).

Bacterial polysaccharide biosynthesis proceeds through distinct biochemical pathways. Xanthan synthesis follows the Wzx/Wzy-dependent mechanism, which generates heteropolymeric structures through sequential assembly of oligosaccharide repeating units onto a diphosphate membrane anchor (Schmid et al., 2015). The chromosomal *gum* gene cluster regulates xanthan biosynthesis in *Xcc*, spans about 16 kilobases, and encodes 12 structural genes (*gumB* - *gumM*). Upstream bifunctional genes *xanA* and *xanB* supply essential nucleotide-sugar precursors (Becker et al., 1998; Schmid et al., 2015). The repeating unit of xanthan consists of two glycosidic β-1,4-linked glucoses, with a side chain on the nonreducing end made from an α-1,3-linked mannose, a β-1,2-linked glucuronic acid, and a β-1,4 linked mannose (Fig. 1). The production of the repeating units starts with the so-called initiating glycosyltransferase GumD, which catalyzes transfer of glucose-1-phosphate from UDP-glucose to the C55 lipid carrier embedded in the cytoplasmic membrane. GumM extends the nascent structure by adding a second glucose via β-1,4-linkage. GumH then incorporates the first mannose residue from GDP-mannose through α-1,3-attachment to every second glucose. GumK transfers glucuronic acid from UDP-D-glucuronate via β-1,2-bond formation. GumI completes pentasaccharide assembly by attaching the terminal mannose through β-1,4-linkage (Becker et al., 1998; Schmid et al., 2015). Mannose residues undergo modification with acetyl or pyruvyl substituents. The acetyltransferases GumF and GumG acetylate the inner α-mannose and the terminal β-mannose positions, respectively, while the pyruvyltransferase GumL provides 4,6-O-pyruvylation for the outer β-mannose (Katzen et al., 1998). The flippase GumJ (Wzx homolog) facilitates translocation of lipid-linked repeating units into the periplasm. Polymerization proceeds through GumE (Wzy protein) activity, with the polysaccharide copolymerase GumC controlling final chain length determination (Islam et al., 2010; Marczak et al., 2013; Schmid et al., 2015; Vorhölter et al., 2008; Whitfield, 2006).

**Figure 1:**
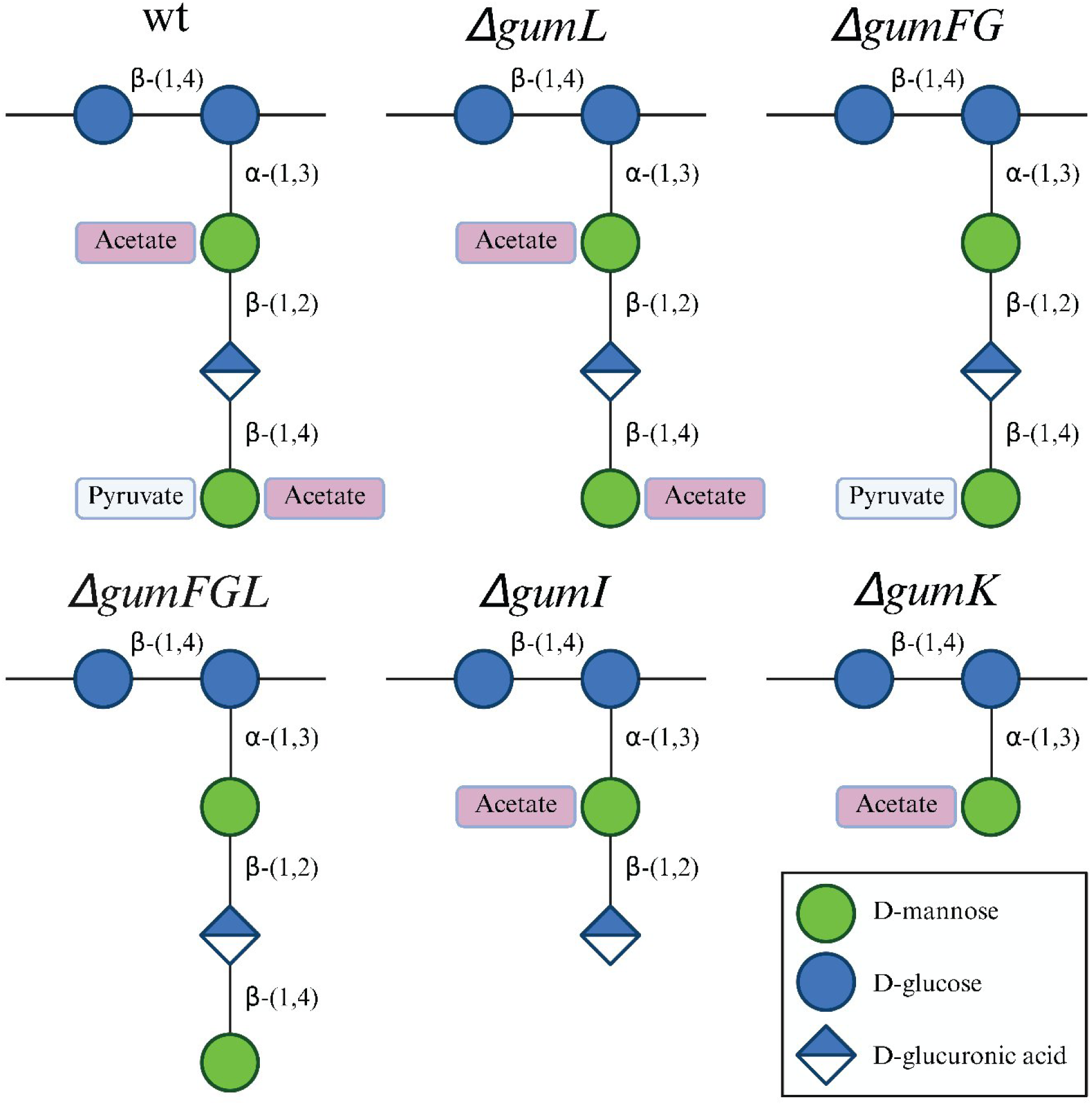
Schematic representation of the xanthan repeating unit structures of the wild type (wt) and the *gum* gene knockout mutants. Deleted genes are indicated in black. Monosaccharides are shown according to the Symbol Nomenclature for Glycans (SNFG): glucose (blue circles), mannose (green circles), and glucuronic acid (blue/white diamond).

Despite extensive genetic insights into xanthan biosynthesis, comprehensive NMR signal assignment and molecular-level structural characterization of the polysaccharide have remained challenging. This limitation arises from intrinsic difficulties associated with the structural analysis of complex polysaccharides, including severe signal overlap, conformational heterogeneity, and restricted spectral resolution due to high molecular weight (Lowicki et al., 2015; Yao et al., 2021). Nuclear magnetic resonance (NMR) spectroscopy provides unparalleled capabilities for polysaccharide structure determination, revealing glycosidic linkage patterns, anomeric configurations, and substitution architectures that remain inaccessible to alternative analytical approaches (Fontana & Widmalm, 2023; Speciale et al., 2022).

However, the structural complexity of xanthan has limited comprehensive resonance assignments using conventional NMR methodologies. Previous investigations have reported only partial assignments based primarily on one-dimensional ^13^C and ^1^H NMR spectra acquired from sonicated and thermally denatured samples, allowing limited identification of characteristic signals such as anomeric carbons and acetyl groups (Cheetham & Mashimba, 1992; Horton et al., 1985). Subsequent NMR studies on xanthan primarily aimed to investigate its physicochemical properties. Xanthan forms an ordered conformation in which two chains interact, forming a duplex conformation (Coviello et al., 1986; Sato et al., 1984). Increasing temperature disrupts inter-chain interactions, leading to the transition to a single-chain disordered state (W. Liu & Norisuye, 1988; Milas & Rinaudo, 1979), although complete denaturation is not achievable in pure water even at 95°C (Hjerde et al., 1994; Kawakami et al., 1990).

Resonance assignments were made only insofar as they were necessary to support these property-focused analyses, rather than to achieve comprehensive structural assignments (Gamini et al., 1991). To date, a complete two-dimensional ^1^H and ^13^C chemical shift assignment of all residues has not been achieved. The challenge is due to the crowding of NMR spectra with overlapping signals from the ring protons (H2–H6). This poses an obstacle to tracing complete spin systems through the signals of five distinct sugar residues.

Targeted gene deletion offers a strategy to overcome these limitations. Strain deletion mutants lacking specific glycosyltransferases or modification enzymes (e.g., Δ*gumK*, Δ*gumI*, Δ*gumF*, Δ*gumG*, Δ*gumL*) generate truncated and modified xanthan polysaccharides (Becker et al., 1998; Doherty et al., 1996; Hassler & Doherty, 1990; Salinas et al., 2011) with simplified NMR spectra, enabling assignment of signals that would typically overlap in wild-type (wt) xanthan spectra (Becker et al., 1998; Salinas et al., 2011). Such genetic approaches have been of use in confirming GumK function as the glucurono-syltransferase and GumI as the terminal mannosyltransferase (Becker et al., 1998).

In this study, we present comprehensive structural characterization of xanthan and biosynthetic variants through an integrated experimental approach combining multi-dimensional NMR spectroscopy, targeted gene knockouts, and functional characterization of the xanthan variants. We achieved near-complete NMR assignments for the pentasaccharide repeating unit through systematic analysis of HSQC, HMBC, TOCSY, and NOESY spectra of xanthan from the wt, complemented by data of samples from the *ΔgumK* and *ΔgumI* knockout strains, and evaluated the weight average molecular weight and order-disorder conformational transitions.

## 2. Materials and methods

### 2.1 Preparation of electrocompetent *X. campestris* cells

Electrocompetent *X. campestris* cells were produced according to the protocol from Wang et al. (Wang et al., 2016) adapted by Gansbiller et al. (Gansbiller et al., 2019). A main culture of *X. campestris* was inoculated into 50 mL growth medium (10 g/L yeast extract, 5 g/L casein peptone, 5 g/L NaCl) from an overnight culture. At an OD_600_ of 0.6 to 1.0, cells were harvested by centrifugation (7,500 g, 4 °C, 10 min) and washed three times with 15 mL 250 mM sucrose solution by resuspending and centrifuging. Finally, cells were resuspended in 2 mL 250 mM sucrose and frozen at −20 °C in 100 µL aliquots until further use.

### 2.2 Creation of *X. campestris* deletion mutants

The deletion mutants of *X. campestris* were created by homologous recombination with the pK19Gm plasmid, a derivative of pK*19mobsacB* from Schäfer et al. (1994) according to Gansbiller et al. (2019). Primers were designed based on the draft genome of *X. campestris* LMG 8031 sequenced by Schmid et al. (2016). For the deletion of genes in *X. campestris*, the regions 500 bp up- and downstream of the gene of interest were amplified by PCR, using isolated genomic DNA of *X. campestris* as a template. These were used as homologous flanks and inserted into the pK19Gm plasmid via isothermal assembly. The assembly mix was used to transform chemically competent *E. coli* Turbo cells by heat shock. Colony PCR and sequencing verified the correctness of the construct.

Electrocompetent *X. campestris* cells were transformed by adding 600-1000 ng of pK19Gm plasmid with the homologous flanks of the respective gene of interest and electroporating at 1,800 V, 400 Ω in a 1 mm cuvette. After recovery with 1 mL Lysogeny Broth (LB) medium (Bertani, 1951) for 2 h at 30 °C, 200 rpm, cells were plated on LB agar containing 30 µg/mL gentamicin and incubated for 48 h at 30 °C. A single colony was inoculated in 3 mL LB medium for 24 h and subsequently plated on LB agar with 10 % (w/v) sucrose to trigger the second homologous recombination. The deletion of the gene of interest was verified by colony PCR and sequencing.

### 2.3 Cultivation of *X. campestris* strains to produce xanthan variants

All xanthan variants were produced in a DASGIP parallel fermenter system in a 600 mL scale equipped with 3 Rushton impellers over 48 h with a controlled pH of 6.8 and a pO_2_ of 30 % saturation. Fermentations were started with an OD_600_ of 0.1 by inoculation of an appropriate volume from an overnight preculture. Cultivations of *X. campestris* and mutants thereof were carried out in EPS medium (5 g/L casein peptone, 1.33 g/L MgSO_4_x7 H_2_O, 0.05 g/L CaCl_2_x2 H_2_O, 1.67 g/L KH_2_PO_4_, 30 g/L glucose, 2 mL/L RPMI 1640 vitamins solution (Sigma-Aldrich) and 1 mL/L trace elements (2.5 g/L iron(II) sulfate heptahydrate, 2.1 g/L sodium tartrate dihydrate, 1.8 g/L manganese(II) chloride tetrahydrate, 0.075 g/L cobalt(II) chloride hexahydrate, 0.031 g/L copper(II) sulfate heptahydrate, 0.258 g/L boric acid, 0.023 g/L sodium molybdate and 0.021 g/L zinc chloride)). To recover xanthan variants from the cultivation broth, the cultures were centrifuged for 30 min at 18,000 x g, 40 °C to remove cells. The supernatant was mixed with isopropanol at a ratio of 1:3 under continuous stirring to precipitate xanthan from the solution. The precipitate was collected and dried at 40 °C, 40 mbar for 24 h in a vacuum oven.

### 2.4 Xanthanase production

The gene encoding the RuGH5a xanthanase was identified by Ostrowski et al. and synthesized by Twist Bioscience in a pET-28(+) vector for *E. coli* expression (Ostrowski et al., 2022), containing a C-terminal His_6_-tag. Chemically competent *E. coli* BL21(DE3) cells were transformed by heat-shock. Individual colonies were cultured in 5 mL of LB medium, supplemented with 50 µg/mL kanamycin at 37 °C and 225 rpm for 6 h, after which these starter cultures were used to inoculate 500 mL of 2×LB medium containing 50 µg/mL kanamycin. Cultures were grown for 24 h at 20 °C in a LEX bioreactor (Epiphyte3) before induction with IPTG at a final concentration of 0.1 mM to activate the promoter and initiate protein expression. RuGH5a was expressed for 24 h, and cells were harvested by centrifugation at 25,000 × g for 20 min at 4 °C. The resulting cell pellet was resuspended in 25 mL of Buffer A (50 mM Tris-HCl, 300 mM NaCl, pH 8.0) and purified using a 1 mL HisTrap™ FF Crude column (Cytiva) on an ÄKTA pure™ system, with Buffer B (50 mM Tris-HCl, 300 mM NaCl, 400 mM imidazole, pH 8.0) for elution (Fig. S11). The eluate was further purified by size-exclusion chromatography using a Superdex® 75 Increase 10/300 GL column (Cytiva) on an ÄKTA pure™ system (Fig. S12). The collected fractions were analyzed by SDS–PAGE on 12% SurePAGE Bis-Tris gels (GenScript) (Fig. S13). Protein bands were compared against the PAGE-MASTER Protein Standard Plus (GenScript) to confirm the presence of RuGH5a (99.1 kDa). Fractions containing RuGH5a were combined and concentrated using a Vivaspin® centrifugal concentrator (50 kDa MWCO; Sartorius). The protein concentration was quantified by measuring absorbance at 280 nm on a NanoDrop ND-1000 spectrophotometer (Thermo Fisher Scientific) using the extinction coefficient (ε = 192435 M⁻¹ cm⁻¹). The purified protein was stored and used in 50 mM MES, 100 mM NaCl, pH 6.0.

### 2.5 Xanthan treatment

The molecular weight of xanthan was reduced by applying shear forces, cycling 50 mg of material in 50 mL deionized water through a Sugino Star Burst wet jet mill homogenizer at 200 MPa for 10 cycles.

Afterwards, the material was dialyzed in 10 L of deionized water and subsequently freeze-dried overnight. 10 mg of the Star Burst treated material was redissolved in 1 mL of phosphate-buffered saline (PBS, pH 6) and subjected to enzymatic degradation using 100 µg of the RuGH5a xanthanase for 4 days at 37 °C at 250 rpm. Afterwards, the sample was freeze-dried overnight and dissolved in D₂O for NMR analysis.

### 2.6 NMR measurements

Most NMR experiments were carried out on Bruker 800 MHz Avance Neo equipped with a 5-mm cryogenic CP-TCI z-gradient probe and SampleJet at the NV-NMR-Center/Norwegian NMR Platform in Trondheim, Norway. Data acquisition and processing were performed using Bruker TopSpin 4.4.0. Spectral assignments were annotated in Inkscape version 1.4.2. All experiments were conducted at 298.1 K. Pulse programs, acquisition settings, and processing parameters were kept identical across all xanthan variants within the NMR experiment types to ensure comparability of the resulting spectra.

#### Resonance assignment

The HSQC spectra were acquired using the Bruker standard pulse program hsqcedetgpsisp2.3 with 8 scans, a spectral size of 2048 × 400 points, and a relaxation delay (D1) of 1.5 s. The coupling constant (CNST2) was set to 145 Hz. The HMBC spectra were recorded using the Bruker standard pulse program hmbcetgpl3nd with 32 scans, a spectral size of 2048 × 400 points, and a D1 of 1.5 s. For HMBC, CNST2 was likewise set to 145 Hz, and CNST13 was set to 8 Hz. The TOCSY spectra were measured using the Bruker standard pulse program dipsi2gpphpr with 8 scans, a spectral size of 2048 × 512 points, D1 of 1.5 s and a mixing time (D9) of 100 ms. The NOESY spectra were measured using the Bruker standard pulse program noesygpphpr with 8 scans, a spectral size of 4096 × 512 points, D1 of 2 s and a D9 of 60 ms. Selective TOCSY spectra were measured using the Bruker standard pulse program seldigpzs with 64 scans, a spectral size of 65536 points, D1 of 2 s and a mixing time (D9) of 80 ms. Selective COSY spectra were measured using the Bruker standard pulse program selcogp with 64 scans, a spectral size of 65536 points and a P12 of 26 ms.

#### Estimation of the degrees of substitution

Acetylation and pyruvylation degrees at the individual side-chain positions were determined by integration of resolved sub-population signals in the 1D ¹H spectrum of the wt sample. Because the 4,6-O-pyruvate ketal and the 6-O acetyl group are mutually exclusive on the terminal β-mannose (Kool et al., 2013), the terminal-mannose population is fully partitioned into three states: pyruvylated (Pyr), acetylated (Ac) and non-substituted (ns). The degree of pyruvylation (DSPyr) was obtained from the integrals (I) of the pyruvate methyl singlet, normalized to three protons per pyruvylated unit; the degree of acetylation of the terminal mannose (DSAc,βMan) from a single downfield-shifted H6 proton of the 6-O-acetylated β-mannose (one proton per unit); and the non-substituted fraction from the anomeric proton (H1) of the β-mannose (one proton per unit).

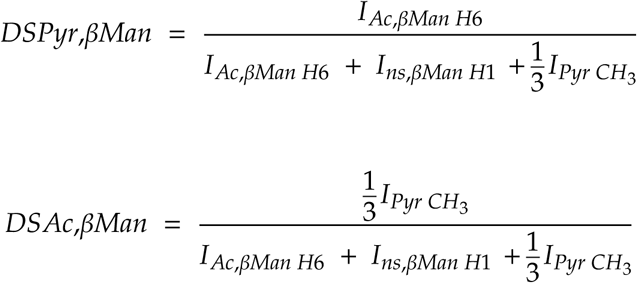

For the inner mannose, the resolved anomeric protons of the acetylated and non-substituted α-mannose (one proton each). Integrals were extracted after phase and baseline correction in Bruker TopSpin 4.4.0.

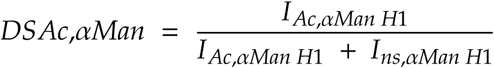

#### Transition temperatures

Rigid structures like the xanthan ordered conformation tumble poorly in solution, leading to rapid relaxation and broad NMR peaks. Heating of xanthan induces an order-disorder transition that increases mobility and sharpens the resonances (Morris et al., 1977). This behavior allows plotting the integral of a suitable reporter peak as a function of temperature. This results in a sigmoidal curve that can be described by a two-state transition. The melting temperature (Tm) is the inflection point of the transition and can be estimated using the Boltzmann equation (Niesen et al., 2007):

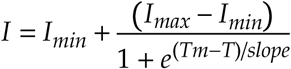

The *Tm* corresponds to the temperature at which 50% of the sample is considered denatured and represents the midpoint of the order–disorder transition. In the sigmoidal fit, *I* denotes the measured intensity, while *I*_max_ and *I*_min_ define the upper and lower asymptotes of the curve. The slope parameter reflects the steepness of the transition. If the dataset could not be adequately fitted with a sigmoidal function, the model was modified to estimate the melting temperature as the midpoint of the transition in a non-sigmoidal function:

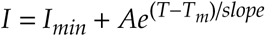

*A* describes the difference between *I*_min_ and *I*_max_. This applies if all temperature points are below the *Tm*. All T_m_ measurements were performed on samples after ten cycles of mechanical treatment (see section 2.5). The spectra used to determine the transition temperatures were acquired using the Bruker standard pulse program zg30 on a Bruker 600 MHz Avance Neo equipped with a 5-mm Smartprobe with z-gradient and SampleCase, at the NV-NMR-Center/Norwegian NMR Platform in Trondheim, Norway. Spectra were acquired at 298 K, 313 K, 318 K, 323 K, 328 K, 333 K, 338 K, 343 K, 348 K, 353 K, 358 K, and 363 K. The spectra were measured using 32 scans, a spectral size of 65536 points, and a D1 of 1.0 s. The data were acquired and processed using Bruker TopSpin 4.4.0. The integrals of the reporter peak at each temperature were fitted to a sigmoidal curve to determine or estimate the melting temperatures of the xanthan variants.

### 2.7 SEC-MALS

Size-Exclusion Chromatography with Multi-Angle Laser Scattering detection (SEC-MALS) was used to monitor the weight average molecular weight (Mw) across successive cycles (0 to 10) of mechanical treatment by a wet jet mill homogenizer. The dried EPS was ground using the Retsch Ultra Centrifugal Mill ZM 200 for 10 s at 18,000 rpm. 1 g/L EPS was dissolved in 10 mL of SEC-MALS buffer (17 g/L NaNO_3_, 8.995 g/L Na_2_HPO4, 0.2 g/L NaN_3_) using a speed mixer for 1 min at 2,500 rpm and subsequent overnight stirring. The samples were then filtered with a 0.45 μm nylon syringe filter.

2 x 300 x 8 mm AppliChrom ABOA SuperOH-P450 columns with a separation range between 50 kDa and 10 MDa and a 50 x 8 mm precolumn was used with a temperature of 38 °C, SEC-MALS buffer as the mobile phase flowing at 0.8 mL/min, and a sample injection volume of 100 μL every 100 min. The PN3621 MALS detector was operated with a LS-power of 80 %, a sampling frequency of 2 Hz. A dn/dc value of 0.147 mL/g and an extinction coefficient of 0.667 mL/g*cm, and the random coil fit type with a fit degree of 1 was used to evaluate the signals. Only scattering angles with an appropriate signal and a defined peak were chosen for analysis. Measurements were performed in technical triplicates for commercial xanthan, Δ*gumI* and Δ*gumK*, and in duplicates for wt, Δ*gumFG*, Δ*gumL* and Δ*gumFGL*.

## 3. Results

### 3.1 Chemical shift assignment

Chemical shift assignments provide a valuable basis for the structural characterization of modified polysaccharides, as they enable the detection of subtle changes in polymer architecture. To establish such assignments for xanthan, we recorded HSQC, HMBC, TOCSY, and NOESY spectra, as well as selective TOCSY and selective COSY spectra of xanthan obtained from the wt alongside five genetically engineered variants (Table 1). Comparing any two variants that differ by exactly one side-chain element identifies the resonances of that element: peaks present in wt but absent in the Δ*gumI* variant belong to the terminal β-mannose and its pyruvyl group, and peaks present in Δ*gumI* but absent in Δ*gumK* belong to glucuronic acid.

**Table 1:**
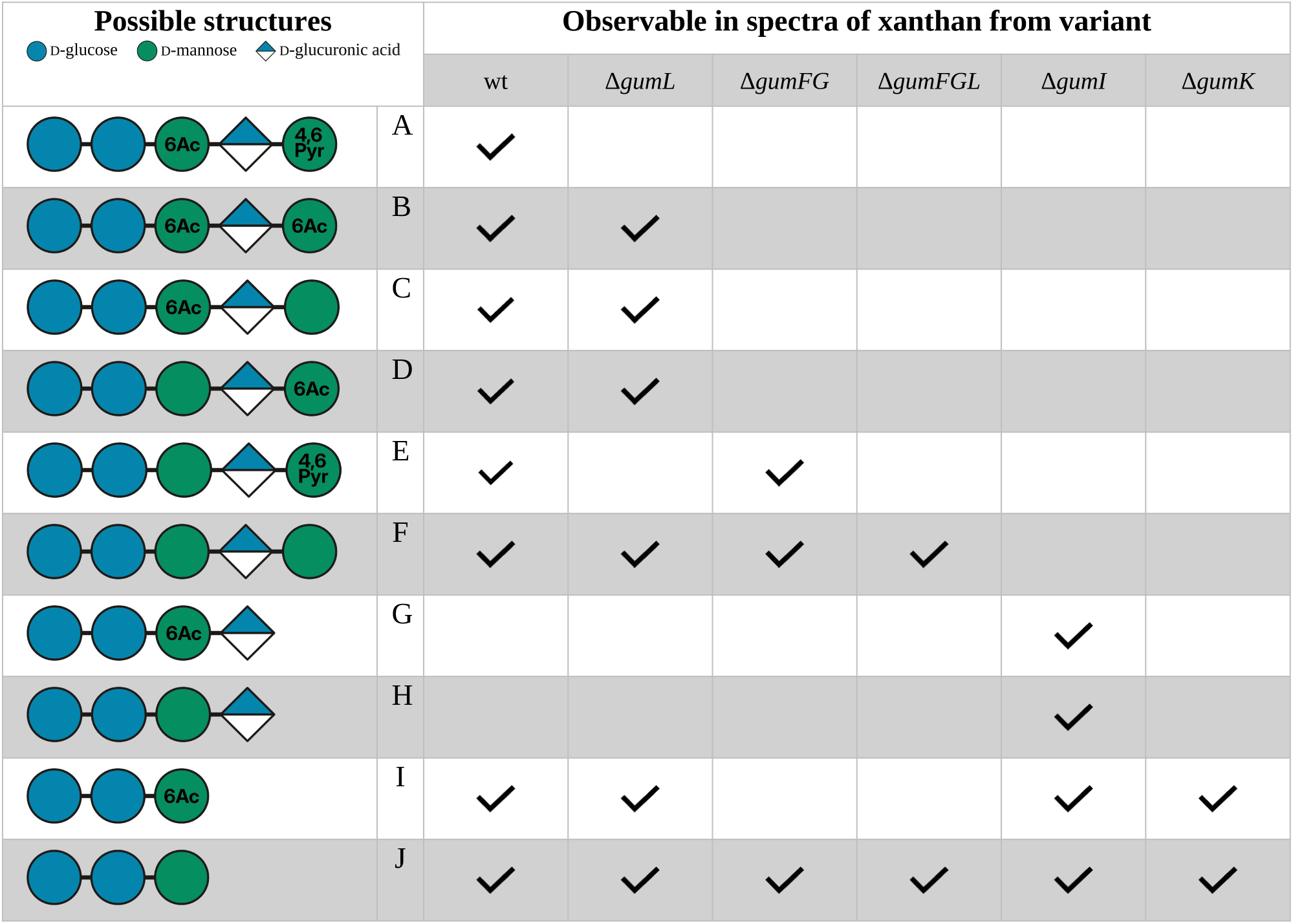
Occurrence of repeating unit structures (Fig. 1) in the spectra of xanthan from the wt and knockout variants. A check mark indicates that the resonances of the structure are present in the spectrum of the respective sample. Because acetylation and pyruvylation are incomplete, a single sample contains several possible structures, and it is not possible to discriminate between overlapping ones such as the α-Man-6Ac signal in A-C. The structures are defined as follows: A: pentasaccharide repeating unit with acetylated α-mannose and pyruvylated β-mannose; B: pentasaccharide repeating unit with acetylated α-mannose and acetylated β-mannose; C: pentasaccharide repeating unit with acetylated α-mannose; D: pentasaccharide repeating unit with acetylated β-mannose; E: pentasaccharide repeating unit with pyruvylated β-mannose; F: pentasaccharide repeating unit with no substitution of the mannoses; G: tetramer with acetylated α-mannose; H: tetramer without substitution; I: trimer with acetylated α-mannose; J: trimer without substitution.

| Possible structures<br>D-glucose D-mannose D-glucuronic acid |  | Observable in spectra of xanthan from variant |  |  |  |  |  |
|---|---|---|---|---|---|---|---|
|  |  | wt | Δ*gumL* | Δ*gumFG* | Δ*gumFGL* | Δ*gumI* | Δ*gumK* |
|  | A | ✓ |  |  |  |  |  |
|  | B | ✓ | ✓ |  |  |  |  |
|  | C | ✓ | ✓ |  |  |  |  |
|  | D | ✓ | ✓ |  |  |  |  |
|  | E | ✓ |  | ✓ |  |  |  |
|  | F | ✓ | ✓ | ✓ | ✓ |  |  |
|  | G |  |  |  |  | ✓ |  |
|  | H |  |  |  |  | ✓ |  |
|  | I | ✓ | ✓ |  |  | ✓ | ✓ |
|  | J | ✓ | ✓ | ✓ | ✓ | ✓ | ✓ |

The high molecular weight and restricted molecular mobility at room temperature (Morris, 2019; Morris et al., 1977; Ndukwe et al., 2024) preclude NMR analysis of unprocessed xanthan, whose signals in NMR spectra are broad beyond detectability. To overcome these challenges, xanthan samples were treated with a combination of mechanical degradation and enzymatic depolymerization using the RuGH5a xanthanase (Ostrowski et al., 2022). The resulting samples yielded NMR spectra with sharp signals at room temperature, allowing us to carry out chemical shift assignments.

The five knockout strains used here each abolish side-chain assembly, acetylation or pyruvylation at a defined step (deleted genes in Fig. 1). If every repeating unit were synthesized homogeneously, one would expect a one-to-one correspondence between genotype and structure. For example, Δ*gumL* would correspond to the unpyruvylated side chains (B, C, and D in Table 1). In practice, however, acetylation and pyruvylation are incomplete, so a single sample contains both the decorated and non-decorated populations of α-mannose and β-mannose. Each sample therefore contains not only the structure assigned to its strain (red labels in Fig. 1) but also the structures lacking one or both substitutions (Table 1). Throughout the assignment, we use this subset principle to attribute the resonances observed in each knockout sample to the structures present in it.

The β-mannose was assigned first, as it represents the outermost residue of the xanthan side chain. Comparison of the wt and the Δ*gumI* variant HSQC spectra revealed a distinct set of absent signals in the Δ*gumI* variant. These missing signals were attributed to β-mannose, consistent with the role of GumI in transferring the β-mannose residue onto the glucuronic acid within the side chain (Becker et al., 1998). Two β-mannose units were identified in total, one of which was pyruvylated: it exhibited a detectable correlation between H6 and the pyruvate ^13^C resonance (δ^1^H: 1.46 ppm; δ^13^C: 24.74 ppm) in the wt HMBC spectrum, consistent with a 4,6-O-pyruvate ketal. The order of the resonances (H1/C1–H6/C6) for pyruvylated β-mannose was established using the wt HMBC spectrum, which revealed connectivities between H1/C2, H3/C4, and H5/C6. NOESY correlations provided cross-peaks for assigning H2 and H4, as well as H1, H3, and H5, clarifying the order of H/C pairs. The assignment was supported by TOCSY data and previously reported β-mannose chemical shifts (Agrawal, 1992). For the non-pyruvylated β-mannose, the wt HMBC spectrum indicated correlations between H1/C2 and H3/C4, H4/C5 and H5/C6. NOESY further showed cross-peaks involving H1, H3, and H5. As for the pyruvylated β-mannose, these assignments were confirmed by TOCSY data and published chemical shifts for β-mannose (Agrawal, 1992). The H4/C4 resonance of pyruvylated β-mannose (δ^1^H: 3.69 ppm; δ^13^C: 80.13 ppm) appears downfield (high carbon ppm) relative to both non-pyruvylated β-mannose (δ1H: 3.79 ppm; δ^13^C: 75.59 ppm) and previously reported β-mannose values, which further indicates pyruvylation of the C4 in this β-mannose. These assignments were further supported by selective TOCSY spectra recorded from the H2 resonances of both the pyruvylated and non-pyruvylated β-mannose residues in the wt spectrum. Although the terminal β-mannose of xanthan is known to carry acetyl substituents, we could only identify signals for the H6/C6 of acetylated β-mannose in the HSQC spectra of xanthan from the wt (δ^1^H: 4.47 ppm; δ^13^C: 63.55 ppm and δ^1^H: 4.26 ppm; δ^13^C: 63.55 ppm). The complete spin system could not be assigned. A recent study showed that GumG acetylates this residue non-specifically at the 2-O, 3-O, 4-O, and 6-O positions (Y. Liu et al., 2026), implying that the pyruvylated β-mannose could in principle also carry 2-O or 3-O acetylation in addition to the 4,6-O-pyruvate ketal. In our spectra, however, only 6-O acetylation could be identified.

The assignment of the glucuronic acid followed a similar strategy. Signals missing in the Δ*gumK* HSQC spectrum relative to the Δ*gumI* construct were attributed to the GlcA residue, consistent with the role of GumK in transferring GlcA onto the α-mannose (Becker et al., 1998). The order of the carbon correlations was determined using the wt HMBC spectrum, in which all correlations from C1 to C5 were observed. TOCSY and NOESY spectra from the wt, together with TOCSY and NOESY data from the Δ*gumI* xanthan and previously reported GlcA assignments (Agrawal, 1992), further supported the assignment. Similar to the pyruvylated β-mannose, the C4 resonance appeared downfield compared to previously reported GlcA chemical shifts for this position, likely due to the glycosidic linkage to β-mannose. Even though the C6 resonance could not be assigned, the remaining assignments were further supported by a selective TOCSY spectrum recorded from the C1 resonance of glucuronic acid in the Δ*gumI* xanthan spectrum.

Due to the α-configuration of the first mannose residue in the side chain, its anomeric proton resonates at a higher chemical shift than the anomeric signals of the β-sugars (Agrawal, 1992). Four signals can be observed in that area in the wt HMBC. In addition, the H6 signal of the acetylated α-mannose shows a correlation to the acetate ^13^C resonance in the wt HMBC spectrum. From these anchor points, HMBC correlations were used to assign H2/C2, H3/C3, H4/C4, and H5/C5. However, no direct H2/C3 correlation was observed for the non-acetylated α-mannose. Selective TOCSY and COSY experiments across different spectra revealed highly similar chemical environments for the acetylated and non-acetylated α-mannose, resulting in similar and even overlapping H3/C3, H4/C4, and H5/C5 resonances. Selective TOCSY experiments further revealed two distinct H2/C2 correlations for both acetylated (δ^1^H: 4.18 ppm; δ^13^C: 70.48 ppm & δ^1^H: 4.21 ppm; δ^13^C: 77.79 ppm) and non-acetylated α-mannose (δ^1^H: 3.96 ppm; δ^13^C: 72.60 ppm & δ^1^H: 4.19 ppm; δ^13^C: 77.60 ppm). These signals can be explained by the presence of two variants of the acetylated and non-acetylated α-mannose, one retaining the remainder of the side chain and one lacking the other residues, resulting from disruption of the α-mannose to β-glucuronic acid glycosidic linkage. This is consistent with the disappearance of the H2/C2 correlations corresponding to α-mannose residues retaining the remainder of the side chain in the spectra of the Δ*gumK* variant. The break of the glycosidic bond also provides an explanation for the four anomeric signals that were described earlier. Furthermore, a difference in the C6 chemical shift is observed between acetylated α-mannose (δ^1^H: 4.37 ppm; δ^13^C: 63.40 ppm) and non-acetylated α-mannose (δ^1^H: 3.84 ppm; δ^13^C: 60.01 ppm). The downfield shift of the H6 resonance in acetylated α-mannose can be attributed to the presence of an acetyl group at C6. TOCSY and NOESY spectra for xanthan from the wt, together with previously reported chemical shifts for α-mannose (Agrawal, 1992), were used to confirm the assignments. The peaks corresponding to the α-mannose residues are observable in all HSQC spectra (F and G in Table 1), including those of the variants Δ*gumI* and Δ*gumK*, consistent with the expected presence of α-mannose in their side chains.

A similar approach was used to assign the two glucose units, one with a side chain and one without. All corresponding peaks were observed in the HSQC spectra of the wt, Δ*gumI*, and Δ*gumK* variants, consistent with the presence of both glucose residues in all possible structures (A-J in Table 1). The glucose residues were first identified using TOCSY data, as their anomeric carbons could be distinguished based on the spectra of the gene deletion variants. The order of the remaining resonances was established using HMBC and NOESY spectra from the wt sample, as the HMBC cross-peaks between H3/C3 and H4/C4 for both glucose residues, between H2/C2 and H3/C3 for the glucose with side chain, and between H4/C4 and H5/C5 for the glucose without side chain were absent. The assignments were further supported by previously reported chemical shifts for glucose (Agrawal, 1992). Notably, the H3/C3 resonance of the side-chain-linked glucose (δ^1^H: 3.62 ppm; δ^13^C: 83.17 ppm) appears downfield relative to both the glucose without side chain (δ^1^H: 3.63 ppm; δ^13^C: 74.38 ppm) and literature values for this position. In contrast, the C4 carbon resonance of the glucose without side chain (δ^1^H: 3.63 ppm; δ^13^C: 79.04 ppm) is shifted downfield compared to the glucose with side chain (δ^1^H: 3.55 ppm; δ^13^C: 69.88 ppm) and previously assigned glucose residues. In the wt spectra, an HMBC correlation was observed between the H3 of the glucose with side chain and the C1 of an α-mannose. Additionally, a NOESY correlation was detected between H4 of glucose 2 and H1 of glucose 1. These assignments were further validated by a selective TOCSY spectrum recorded from the C1 and C2 resonances of both glucose residues in the spectrum of the Δ*gumK* variant.

Several additional signals were observed in the wt HSQC spectra that could not be assigned to the xanthan structure and were therefore excluded from the peak assignments (“unknown” in Fig. 2). To distinguish signals originating from xanthan from those introduced during sample preparation, a negative control containing the reaction components (see section 2.5) but no xanthan was subjected to the same incubation and sample processing conditions and analyzed by NMR. Many of the signals labeled “unknown” in Fig. 2, particularly those in the region between δ^1^H: 2.74–3.23 ppm and δ^13^C: 46–53 ppm, were also detected in the negative control, indicating that these signals originate from components of the enzymatic reaction rather than from xanthan. The remaining unassigned signals, which were absent from the negative control, may originate from residual sucrose in the EPS growth medium. Several of these signals exhibit chemical shifts characteristic of glucose and fructose residues that are not part of the xanthan structure (such as the glucose H6/C6 and fructose H1/C1 correlations). These compounds may co-precipitate with xanthan during the isopropanol precipitation step and remain in the sample despite subsequent purification steps.

**Figure 2:**
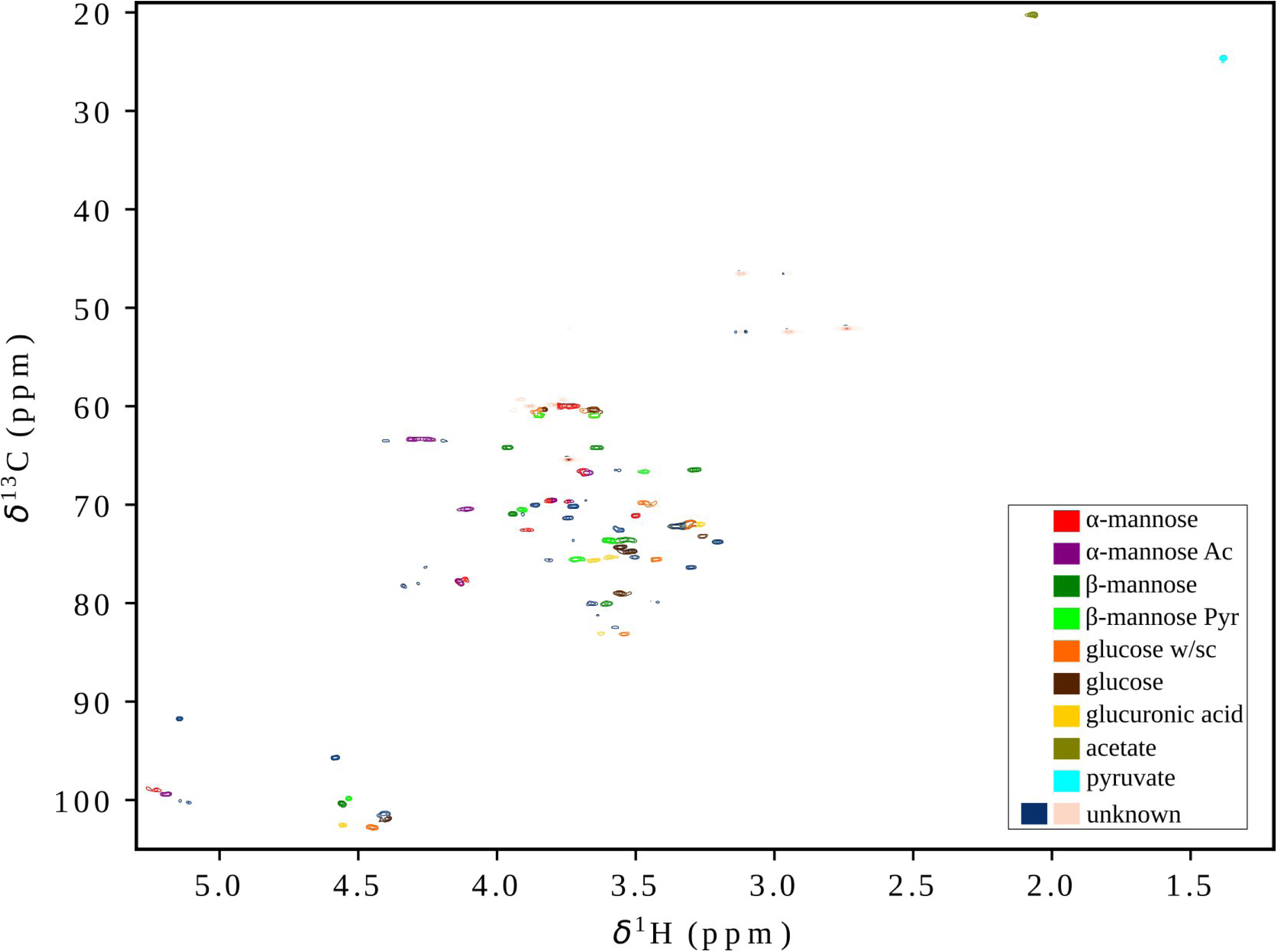
^1^H,^13^C-HSQC chemical shift assignment of xanthan with assigned cross-peaks corresponding to the different sugar residues and their respective carbon–proton pairs. The signal assignments were based on published xanthan NMR data and additional spectra of the xanthan from wt. Red: α-D-mannose without substitution; Purple: acetylated α-D-mannose; Dark green: pyruvylated β-D-mannose; Light green: undecorated β-D-mannose; Orange: β-D-glucose linked to the trisaccharide side chain; Brown: β-D-glucose without a side chain; Yellow: β-D-glucuronic acid; Olive: acetate; Cyan: pyruvate; Dark blue/Peach: unassigned signals (additional peaks not attributable to xanthan).

Figure 2 shows the assigned HSQC spectrum of xanthan from the wt; expanded views of the individual spin systems and a magnified overview are provided in Fig. S2–S9 and the chemical shifts are summarized in Table 2. The wt spectrum serves as the reference because it contains every resonance of the variant spectra, as outlined above. Because the structures share residues, the spectra of different variants also share peaks; Table 1 summarizes, based on our assignment, which structure contributes to each sample and therefore which spectra overlap. Even though the structure J was only expected to be present in the Δ*gumK* spectra, we in fact observed this side-chain-lacking structure in all samples. We propose an explanation for this in the Discussion (section 4.1). This trisaccharide repeating unit can be recognized by the characteristic upfield shift of the α-Man C2 resonance resulting from the absence of GlcA. A comparison of the HSQC spectra obtained for the wt, and all variant constructs is provided in the supplementary data (Fig. S10).

**Table 2:**
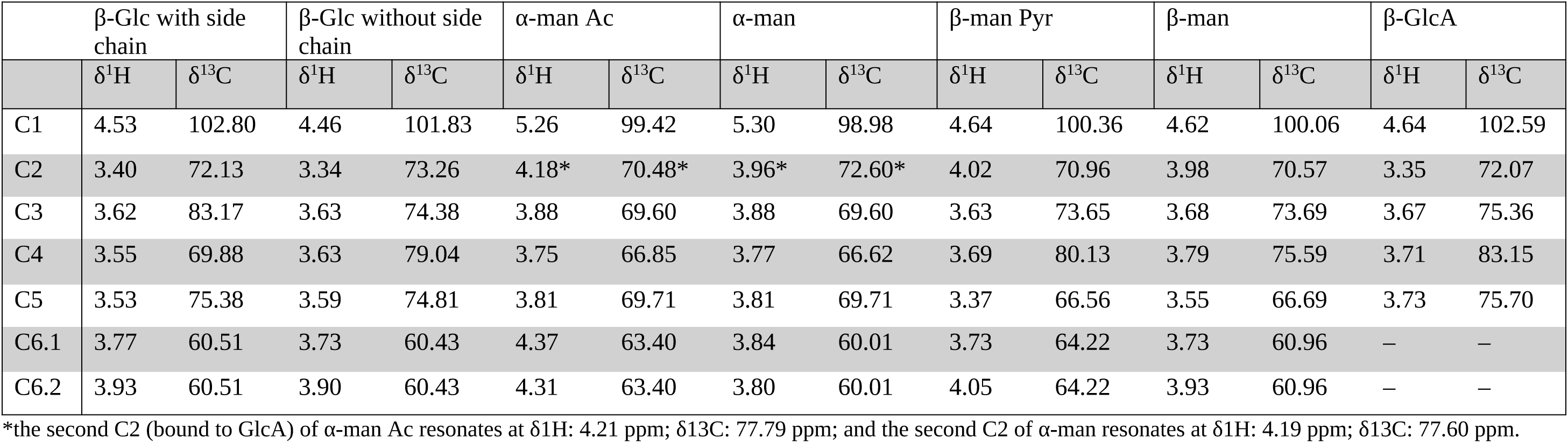
^1^H ^13^C NMR chemical shift assignments (TSP-corrected, ppm) for the individual monosaccharide residues of xanthan. Reported are the proton and carbon chemical shifts for each carbon position (C1–C6) of the respective sugar units. The C6 of the β-glucuronic acid could not be detected. All values are given in ppm.

|  | β-Glc with side chain |  | β-Glc without side chain |  | α-man Ac |  | α-man |  | β-man Pyr |  | β-man |  | β-GlcA |  |
|---|---|---|---|---|---|---|---|---|---|---|---|---|---|---|
|  | δ<sup>1</sup>H | δ<sup>13</sup>C | δ<sup>1</sup>H | δ<sup>13</sup>C | δ<sup>1</sup>H | δ<sup>13</sup>C | δ<sup>1</sup>H | δ<sup>13</sup>C | δ<sup>1</sup>H | δ<sup>13</sup>C | δ<sup>1</sup>H | δ<sup>13</sup>C | δ<sup>1</sup>H | δ<sup>13</sup>C |
| C1 | 4.53 | 102.80 | 4.46 | 101.83 | 5.26 | 99.42 | 5.30 | 98.98 | 4.64 | 100.36 | 4.62 | 100.06 | 4.64 | 102.59 |
| C2 | 3.40 | 72.13 | 3.34 | 73.26 | 4.18* | 70.48* | 3.96* | 72.60* | 4.02 | 70.96 | 3.98 | 70.57 | 3.35 | 72.07 |
| C3 | 3.62 | 83.17 | 3.63 | 74.38 | 3.88 | 69.60 | 3.88 | 69.60 | 3.63 | 73.65 | 3.68 | 73.69 | 3.67 | 75.36 |
| C4 | 3.55 | 69.88 | 3.63 | 79.04 | 3.75 | 66.85 | 3.77 | 66.62 | 3.69 | 80.13 | 3.79 | 75.59 | 3.71 | 83.15 |
| C5 | 3.53 | 75.38 | 3.59 | 74.81 | 3.81 | 69.71 | 3.81 | 69.71 | 3.37 | 66.56 | 3.55 | 66.69 | 3.73 | 75.70 |
| C6.1 | 3.77 | 60.51 | 3.73 | 60.43 | 4.37 | 63.40 | 3.84 | 60.01 | 3.73 | 64.22 | 3.73 | 60.96 | – | – |
| C6.2 | 3.93 | 60.51 | 3.90 | 60.43 | 4.31 | 63.40 | 3.80 | 60.01 | 4.05 | 64.22 | 3.93 | 60.96 | – | – |

\*the second C2 (bound to GlcA) of α-man Ac resonates at δ1H: 4.21 ppm; δ13C: 77.79 ppm; and the second C2 of α-man resonates at δ1H: 4.19 ppm; δ13C: 77.60 ppm.

Integration of the resolved side-chain resonances in the wt sample gave degrees of substitution of 0.77 for 6-O acetylation of the inner α-mannose, 0.68 for 4,6-O-pyruvylation of the terminal β-mannose, and 0.13 for 6-O acetylation of the terminal β-mannose.

### 3.2 Order-disorder transitions

The functional properties of xanthan solutions stem largely from a thermally reversible conformational transition between an ordered, rigid helical structure at low temperature and a disordered, flexible coil at high temperature (Christensen et al., 1993; W. Liu & Norisuye, 1988). The order-disorder transition temperature (Tₘ) is highly sensitive to ionic strength and, critically, to the substitution pattern of the side chains: the pyruvate and acetyl groups and the charged glucuronic acid residues are known to contribute to helix stability (Callet et al., 1987; Dentini et al., 1984; Foss et al., 1987; Shatwell et al., 1990; Smith et al., 1981). Herein, Tₘ values were determined by recording ^1^H NMR spectra from 25 to 90 °C and fitting the integrals of reporter signals to a sigmoidal function, with the midpoint defined as T_m_ (Fig. S1).

Xanthan from wt showed three suitable reporter signals: α-Man H1, the methyl protons of acetate, and the methyl protons of pyruvate. The same approach was applied to the xanthan variants obtained from Δ*gumL* (no pyruvate), Δ*gumFG* (no acetyl), Δ*gumFGL* (no acetyl or pyruvate), Δ*gumK* (no GlcA/β-Man), and Δ*gumI* (no β-Man). For samples with multiple reporters, T_m_ values were generally consistent across reporters, with differences up to ∼10 °C. Using the pyruvate CH_3_ as a reporter gave the lowest T_m_, whereas using the α-Man H1 gave the highest.

For Δ*gumK* and Δ*gumI*, T_m_ could only be determined from the acetate CH_3_, as spectra were recorded in 90% H_2_O/10% D_2_O, causing other signals to overlap with the water resonance. Foss et al. used pure D_2_O for NMR-based and H_2_O for optical rotation-based T_m_ determination of xanthan and noted that a possible isotopic contribution from the solvent could not be separated from the difference between the two methods (Foss et al., 1987); the isotope effect on T_m_ thus remains unquantified. Since both Δ*gumK* and Δ*gumI* are compared against samples measured under the same conditions as wt, the within-series comparison is unaffected, but their absolute T_m_ values should be interpreted with this caveat. Overall, decreasing the pyruvate content (Δ*gumL*, Δ*gumFGL*) or the negative charge content (Δ*gumK*) increased the transition temperatures by 12–17 °C compared with xanthan from the wt, whereas decreasing the acetate content (Δ*gumFG*), and deletion of the terminal β-Man (Δ*gumI*) decreased the stability by 4–7 °C and 12 °C, respectively. Table 3 summarizes all T_m_ values derived from anomeric and methyl reporter signals.

**Table 3:** Order-disorder mid-point transition values or melting temperatures (T_m_) of all the samples based on both the anomeric proton of α-mannose and methyl peaks of the substitutions. Only wt, Δ*gumL*, Δ*gumK*, and Δ*gumI* had an acetate methyl peak, while only wt and Δ*gumFG* had a pyruvate methyl peak.

|  | **α-Man H1 T<sub>m</sub> (°C)** | **Acetate -CH<sub>3</sub> T<sub>m</sub> (°C)** | **Pyruvate -CH<sub>3</sub> T<sub>m</sub> (°C)** |
|---|---|---|---|
| wt | 68.0 ± 1.2 | 62.9 ± 0.4 | 60.6 ± 0.7 |
| Δ*gumL* | ≥85 | ≥ 85 | - |
| Δ*gumFG* | 64.1 ± 1.7 | - | 53.5 ± 0.6 |
| Δ*gumFGL* | 81.1 ± 1.0 | - | - |
| Δ*gumK* | - | 75 ± 4* | - |
| Δ*gumI* | - | 51 ± 2* | - |

\*These samples were measured in 90% H<sub>2</sub>O/10% D<sub>2</sub>O. The acetate methyl proton signal is the only reporter not overlapping with the water signal.

The order-disorder transition is sensitive to molecular weight and aggregation state, and the wt and side-chain variants differed substantially in native molecular weight. A direct comparison of the untreated samples would therefore not have allowed differences in transition temperatures to be attributed to side-chain composition rather than chain length. SEC-MALS was used to quantify how mechanical degradation alters the weight average molecular weight (Mw) distribution of xanthan from the wt and side-chain variants, and to define a common molecular-weight distribution for downstream analyses.

Measurements before mechanical degradation (Fig. 3) yielded very high Mw values (1.8–12.6 MDa). Mechanical treatment with a wet jet mill homogenizer caused a pronounced decrease in Mw during the initial cycles, followed by progressively smaller changes, and after ten cycles all samples converged to Mw values of approximately 3.3–8.5 × 10Z kDa. Mechanical degradation could, in principle, also affect the chemical composition of the polysaccharide; however, in xanthan treated by dynamic high-pressure homogenization, the acetate and pyruvate contents remained constant throughout the treatment, indicating that the side chains were not affected (Laneuville et al., 2013).

**Figure 3:**
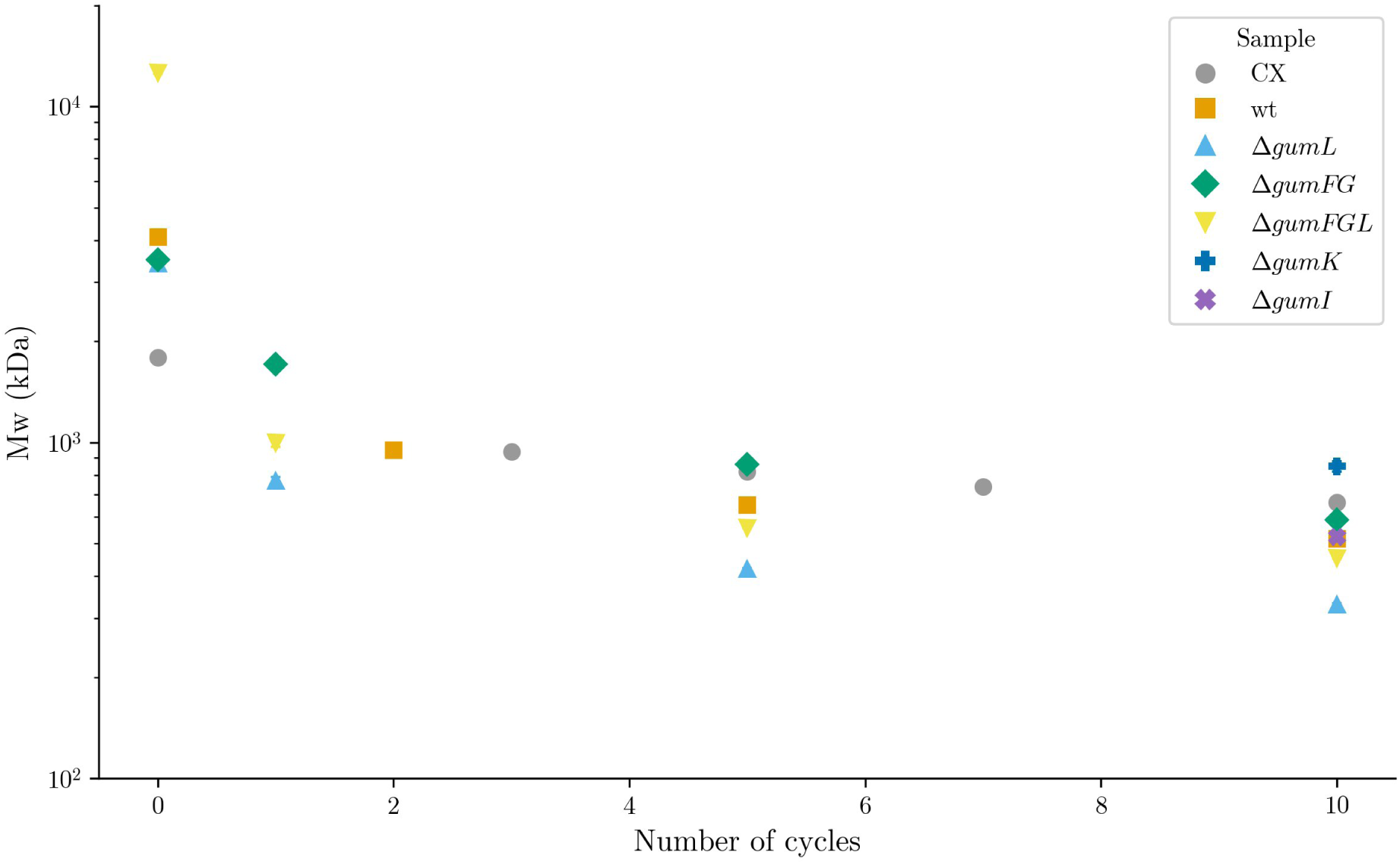
Weight average molecular weight (Mw) for commercial xanthan (CX), wt and variant samples, measured over successive cycles of mechanical treatment with a wet jet mill homogenizer. CX: cycles 0, 3, 5, 7, 10; wt: cycles 0, 2, 5, 10; Δ*gumL:* cycles 0, 1, 5, 10; Δ*gumFG:* cycles 0, 1, 5, 10; Δ*gumFGL:* cycles 0, 1, 5, 10; Δ*gumK:* cycle 10; Δ*gumI*: cycle 10. Error bars represent standard deviations from n=3 for CX, Δ*gumK,* Δ*gumI* and n=2 for wt, Δ*gumL,* Δ*gumFG* and Δ*gumFGL*; for most data points they are smaller than the marker symbols.

## 4. Discussion

As outlined in the Introduction, the high molecular weight of xanthan, its ordered conformation, and severe spectral overlap have so far precluded a complete resonance assignment of the repeating unit. The degradation strategy and variant production described above remove these obstacles, and the resulting assignment is the basis for the functional characterization of engineered variants.

### 4.1 Enzymatic breakdown and genetically engineered xanthan variants enabled thorough NMR characterization

To overcome the challenges outlined above, a combined mechanical and enzymatic degradation strategy was employed. First, xanthan samples were homogenized using a wet jet mill, which reduced polymer length while preserving the ordered conformation. Subsequently, the RuGH5a xanthanase was used to depolymerize the material into repeating-unit oligosaccharides, thereby eliminating the ordered state and substantially supporting the recording of the different spectra. To simplify the resonance assignment of the complex spectra, Δ*gumI* and Δ*gumK* xanthan variants were analyzed using the same workflow. The absence of specific side-chain residues in these variants resulted in less crowded spectra and further enabled the identification of signals originating from β-mannose and β-glucuronic acid residues, when compared to the wt spectra. This reduction in spectral complexity helped to assign the remaining resonances in xanthan from the wt.

The NMR assignment of the xanthan repeating unit revealed pronounced downfield carbon chemical shift changes at the H3/C3 position of one glucose residue and at the H4/C4 position of the glucuronic acid residue. These deshielding effects can be attributed to the formation of glycosidic linkages within the repeating unit, as the observed Δδ^13^C values fall within the characteristic range (+4–11 ppm) typically reported for glycosylation-induced shifts in polysaccharides (Fontana & Widmalm, 2023; Shashkov et al., 1988; Söderman et al., 1998). Notably, only every second glucose unit in the xanthan backbone is substituted at the C3 position with the trisaccharide side chain, whereas the glucuronic acid residue forms a glycosidic linkage at C4 with the β-mannose unit, explaining the shift changes (Patel et al., 2020).

Additionally, the H4/C4 correlation of the substituted glucose was shifted upfield relative to that of the unsubstituted glucose. This can be explained by the cleavage specificity of the RuGH5a enzyme, which hydrolyzes the β-(1→4) linkage at the reducing end of the non-branching glucose (Ostrowski et al., 2022). Consequently, the C4 position of the branched glucose is no longer involved in a glycosidic linkage, resulting in increased shielding and therefore an upfield shift compared to the C4 of non-branched glucose, which remains glycosidically substituted.

Furthermore, the NMR data revealed a pronounced downfield shift of the H4/C4 resonance of the pyruvylated β-mannose relative to non-decorated β-mannose and to previously reported literature values. This deshielding effect is consistent with pyruvylation at the C4 position of the pyruvylated β-mannose (Patel et al., 2020). The introduction of a pyruvate substituent to other polymers has been shown to significantly influence the carbon chemical shift due to its electron-withdrawing character (Agnihotri & Misra, 2006). The C6 resonance of the glucuronic acid residue could not be assigned in the recorded spectra. This signal is expected to appear in the downfield region at approximately 176 ppm (Agrawal, 1992). However, the spectral window employed in the present measurements was limited to 105 ppm in the C dimension and therefore did not include the region in which the C6 signal would be expected. Additionally, the acetylated β-mannose residue could not be fully assigned in the recorded spectra, as only the C6/H6 could be identified due to its HMBC correlation with the acetate and the absence of the C6/H6 peaks in the Δ*gumI* spectra. This is likely due to its comparatively low abundance, as we found acetylation only on 13% of the β-mannose units (see section 3.1), whereas pyruvylated and non-mod-ified β-mannose residues were present at higher relative proportions, consistent with previous reports (Kool et al., 2013). Consequently, the corresponding resonances are expected to exhibit reduced signal intensity and may be obscured by overlapping signals from the more abundant structural variants, thereby preventing the identification under the applied experimental conditions. However, the C6/H6 resonances occur in a region with minimal signal overlap and were therefore distinguishable. This allowed the identification of the possible structures containing acetylated β-mannose in the spectra from the different variants (B and D in Table 1).

The occurrence of two α-mannose H2/C2 correlations can be explained by cleavage of the axial glycosidic linkage between C2 of α-mannose and C1 of glucuronic acid during the mechanical treatment with a wet jet mill homogenizer, which exposes the sample to increased temperatures. These observations may be related to the axial configuration of the glycosidic linkage, as axial glycosidic bonds have higher hydrolytic reactivity (Jensen & Bols, 2003) and are more heat labile (Daskhan & Jayaraman, 2011) than their equatorial counterparts. This break also explains the presence of four α-mannose H1/C1 correlations, as cleavage of the linkage results in two populations of both acetylated and non-acetylated α-mannose, with and without the GlcA and β-mannose. However, no second GlcA H1/C1 resonance was observed, as the dialysis performed after the mechanical treatment likely removed any possible GlcA–β-Man disaccharide generated during the treatment. The cleavage of the glycosidic bond also provides an explanation for why structure J (Table 1) is observable in the spectra of all variants, contrary to our expectations, as it can occur across all variants during mechanical treatment. Structure I can also be observed in all spectra except those of Δ*gumFG* and Δ*gumFGL* (Table 1), in which acetylation is abolished. The difference in chemical shifts observed for C6 and H6 of the acetylated and non-acetylated mannose (Table 2) can be attributed to 6-O-acetylation. Addition of the acetyl group induces characteristic chemical-shift changes at the substituted position, resulting in a downfield displacement of both the carbon and proton directly attached to the acetylated carbon (Torzewska et al., 2001).

### 4.2 Side-chain contributions to the thermal stability of the ordered state

The order-disorder transition in xanthan is a reversible, cooperative conformational change in which ordered duplexes (Morris, 2019; Sato et al., 1984) convert to disordered chains upon heating, and reform on cooling, with the transition temperature and kinetics strongly influenced by both solution conditions and side-chain substitution. The acetate and pyruvate content of these genetically engineered xanthan variants has been reported previously (Gansbiller et al., 2019). Our results show that variants lacking pyruvylation (Δ*gumL*, Δ*gumFGL*) exhibit higher conformational transition temperatures (Table 3), consistent with depyruvylated xanthan having been shown to have higher transition temperatures (Callet et al., 1987; Christensen et al., 1993; Dentini et al., 1984; Shatwell et al., 1990; Tomofuji et al., 2022; Wu et al., 2021). This is consistent with observations that increasing ionic strength stabilizes the ordered conformation and raises T_m_ (Foss et al., 1987; Holzwarth & Ogletree, 1979; W. Liu & Norisuye, 1988; Norton et al., 1984), implying that reducing the effect of electrostatic repulsion between negatively charged pyruvate groups, either by screening or by removing pyruvate, stabilizes the ordered conformation. This also explains the stabilizing effect of the Δ*gumK* variant, which lacks not only pyruvate but also the β-mannose and glucuronic acid, thus further removing the charge density and stabilizing interchain associations in the ordered state.

Following the logic of the previous paragraph, one would expect that Δ*gumI*, which lacks β-mannose and thus also pyruvate, would be more stable than the wt. However, our results show a clear destabilization. Christensen et al. found that partial hydrolysis of the β-mannose preserves the basic order-disorder transition, but alters the magnitude, indicating that interactions with β-mannose contribute substantially to the transition enthalpy (Christensen et al., 1993). This suggests that complete removal of β-mannose eliminates key enthalpic stabilization, for example, hydrogen-bonding interactions in the ordered conformation, and thereby lowers the transition temperature. Notably, X-ray fiber diffraction of a terminal-mannose-less variant showed an essentially unchanged c-repeat and helix symmetry, indicating that the ordered backbone conformation is retained with a reduced molecular diameter (Millane & Narasaiah, 1990).

Removal of acetate (Δ*gumFG*) reduced T_m_ (Table 3), consistent with earlier chemical-modification studies (Dentini et al., 1984; Callet et al., 1987). In the ordered conformation, the side chains pack against the backbone through a network of potential intramolecular hydrogen bonds (Moorhouse et al., 1977), and the acetyl groups on the inner α-mannose (Pelletier et al., 2001; Tako & Nakamura, 1984) or outer β-mannose (Kool et al., 2014) have been proposed to contribute to this stabilization The Δ*gum-FGL* variant, which lacks both acetate and pyruvate, exhibited a higher T_m_ than wt but lower than Δ*gumL* (pyruvate-free, acetate-intact), establishing that the stabilizing contribution of pyruvate removal dominates over the destabilizing contribution of acetate removal (Table 3).

## 5. Declaration of generative AI and AI-assisted technologies in the manuscript preparation process

During the preparation of this work, the authors used GPT NTNU to copy edit the manuscript. After using this tool, the authors reviewed and edited the content as needed and take full responsibility for the content of the published article.

## 6. Data statement

Data and scripts used for analysis and for generating the figures are available from https://github.com/gcourtade/2026_Wegner_xanthan-nmr

## 7. Acknowledgements

This work was supported by the Novo Nordisk Foundation (grant number NNF22OC0073963 to G.C.) and the Norwegian NMR Platform (NNP; grant number 226244 from the Research Council of Norway. The authors thank Dr. Olav A. Aarstad for his invaluable technical assistance with SEC-MALS.

## References

Agnihotri, G., & Misra, A. K. (2006). Efficient synthesis of pyruvate ketals of carbohydrates. Tetrahedron Letters, 47(48), 8493–8497. 10.1016/j.tetlet.2006.09.159

Agrawal, P. K. (1992). NMR Spectroscopy in the structural elucidation of oligosaccharides and glycosides. Phytochemistry, 31(10), 3307–3330. 10.1016/0031-9422(92)83678-R

Becker, A., Katzen, F., Pühler, A., & Ielpi, L. (1998). Xanthan gum biosynthesis and application: A biochemical/genetic perspective. Applied Microbiology and Biotechnology, 50(2), 145–152. 10.1007/s002530051269

Bertani, G. (1951). STUDIES ON LYSOGENESIS I: The Mode of Phage Liberation by Lysogenic Escherichia coli. Journal of Bacteriology, 62(3), 293–300. 10.1128/jb.62.3.293-300.1951

Callet, F., Milas, M., & Rinaudo, M. (1987). Influence of acetyl and pyruvate contents on rheological properties of xanthan in dilute solution. International Journal of Biological Macromolecules, 9(5), 291–293. 10.1016/0141-8130(87)90068-7

Cheetham, N. W. H., & Mashimba, E. N. M. (1992). Proton and carbon-13 NMR studies on xanthan derivatives. Carbohydrate Polymers, 17(2), 127–136. 10.1016/0144-8617(92)90106-Z

Christensen, B. E., Knudsen, K. D., Smidsrød, O., Kitamura, S., & Takeo, K. (1993). Temperature-Induced conformational transition in xanthans with partially hydrolyzed side chains. Biopolymers, 33(1), 151–161. 10.1002/bip.360330115

Coviello, T., Kajiwara, K., Burchard, W., Dentini, M., & Crescenzi, V. (1986). Solution properties of xanthan. 1. Dynamic and static light scattering from native and modified xanthans in dilute solutions. Macromolecules, 19(11), 2826–2831. 10.1021/ma00165a027

Daskhan, G. C., & Jayaraman, N. (2011). Increased glycosidic bond stabilities in 4-*C*-hydroxymethyl linked disaccharides. Carbohydrate Research, 346(15), 2394–2400. 10.1016/j.carres.2011.08.030

Dentini, M., Crescenzi, V., & Blasi, D. (1984). Conformational properties of xanthan derivatives in dilute aqueous solution. International Journal of Biological Macromolecules, 6(2), 93–98. 10.1016/0141-8130(84)90070-9

Doherty, D. H., Ferber, D. M., Marrelli, J. D., Vanderslice, R. W., & Hassler, R. A. (1996). Genetic control of acetylation and pyruvylation of xanthan based polysaccharide polymers (United States Patent No. US5514791A). https://patents.google.com/patent/US5514791A/en

Fontana, C., & Widmalm, G. (2023). Primary Structure of Glycans by NMR Spectroscopy. Chemical Reviews, 123(3), 1040–1102. 10.1021/acs.chemrev.2c00580

Foss, P., Stokke, B. T., & Smidsrød, O. (1987). Thermal stability and chain conformational studies of xanthan at different ionic strengths. Carbohydrate Polymers, 7(6), 421–433. 10.1016/0144-8617(87)90010-5

Gamini, A., De Bleijser, J., & Leyte, J. C. (1991). Physico-chemical properties of aqueous solutions of xanthan: An n.m.r. study. Carbohydrate Research, 220, 33–47. 10.1016/0008-6215(91)80004-7

Gansbiller, M., Schmid, J., & Sieber, V. (2019). In-depth rheological characterization of genetically modified xanthan-variants. Carbohydrate Polymers, 213, 236–246. 10.1016/j.carbpol.2019.02.055

Hassler, R. A., & Doherty, D. H. (1990). Genetic Engineering of Polysaccharide Structure: Production of Variants of Xanthan Gum in Xanthomonas campestris. Biotechnology Progress, 6(3), 182–187. 10.1021/bp00003a003

Hjerde, T., Kristiansen, T. S., Stokke, B. T., Smidsrød, O., & Christensen, B. E. (1994). Conformation dependent depolymerisation kinetics of polysaccharides studied by viscosity measurements. Carbohydrate Polymers, 24(4), 265–275. 10.1016/0144-8617(94)90070-1

Holzwarth, G., & Ogletree, J. (1979). Pyruvate-free xanthan. Carbohydrate Research, 76(1), 277–280. 10.1016/0008-6215(79)80030-0

Horton, D., Mols, O., Walaszek, Z., & Wernau, W. C. (1985). Structural and biosynthetic studies on xanthan by 13C-n.m.r. Spectroscopy. Carbohydrate Research, 141(2), 340–346. 10.1016/S0008-6215(00)90467-1

Islam, S. T., Taylor, V. L., Qi, M., & Lam, J. S. (2010). Membrane Topology Mapping of the O-Antigen Flippase (Wzx), Polymerase (Wzy), and Ligase (WaaL) from Pseudomonas aeruginosa PAO1 Reveals Novel Domain Architectures. mBio, 1(3), e00189–10. 10.1128/mBio.00189-10

Jensen, H. H., & Bols, M. (2003). Steric Effects Are Not the Cause of the Rate Difference in Hydrolysis of Stereoisomeric Glycosides. Organic Letters, 5(19), 3419–3421. 10.1021/ol030081e

Katzen, F., Ferreiro, D. U., Oddo, C. G., Ielmini, M. V., Becker, A., Pühler, A., & Ielpi, L. (1998). Xanthomonas campestris pv. campestris gum mutants: Effects on xanthan biosynthesis and plant virulence. Journal of Bacteriology, 180(7), 1607–1617. 10.1128/JB.180.7.1607-1617.1998

Kawakami, K., Okabe, Y., & Norisuye, T. (1990). Dissociation of dimerized xanthan in aqueous solution. Carbohydrate Polymers, 14(2), 189–203. 10.1016/0144-8617(90)90030-V

Kool, M. M., Gruppen, H., Sworn, G., & Schols, H. A. (2013). Comparison of xanthans by the relative abundance of its six constituent repeating units. Carbohydrate Polymers, 98(1), 914–921. 10.1016/j.carbpol.2013.07.003

Kool, M. M., Gruppen, H., Sworn, G., & Schols, H. A. (2014). The influence of the six constituent xanthan repeating units on the order–disorder transition of xanthan. Carbohydrate Polymers, 104, 94–100. 10.1016/j.carbpol.2013.12.073

Laneuville, S. I., Turgeon, S. L., & Paquin, P. (2013). Changes in the physical properties of xanthan gum induced by a dynamic high-pressure treatment. Carbohydrate Polymers, 92(2), 2327–2336. 10.1016/j.carbpol.2012.11.077

Liu, W., & Norisuye, T. (1988). Thermally induced conformation change of xanthan: Interpretation of viscosity behaviour in 0.01 m aqueous sodium chloride. International Journal of Biological Macromolecules, 10(1), 44–50. 10.1016/0141-8130(88)90066-9

Liu, Y., Rühmann, B., Melse, O., Bayaraa, T., Kampl, L., Döring, M., & Sieber, V. (2026). Unraveling Xanthomonas acetyltransferase GumG: Decoding catalytic promiscuity, understanding the mechanism, and enhancing enzymatic versatility (p. 2026.06.22.733816). bioRxiv. 10.64898/2026.06.22.733816

Lowicki, D., Buda, S., & Mlynarski, J. (2015). NMR of carbohydrates. In Nuclear Magnetic Resonance (Vol. 44, pp. 407–430). Royal Society of Chemistry. 10.1039/9781782622758-00407

Marczak, M., Dźwierzyńska, M., & Skorupska, A. (2013). Homo- and heterotypic interactions between Pss proteins involved in the exopolysaccharide transport system in *Rhizobium leguminosarum* bv. *Trifolii*. Biological Chemistry, 394(4), 541–559. 10.1515/hsz-2012-0161

Milas, M., & Rinaudo, M. (1979). Conformational investigation on the bacterial polysaccharide xanthan. Carbohydrate Research, 76(1), 189–196. 10.1016/0008-6215(79)80017-8

Millane, R. P., & Narasaiah, T. V. (1990). X-ray fiber diffraction studies of a variant of xanthan gum in which the sidechain terminal mannose unit is absent. Carbohydrate Polymers, 12(3), 315–321. 10.1016/0144-8617(90)90072-Z

Moorhouse, R., Walkinshaw, M. D., & Arnott, S. (1977). Xanthan Gum—Molecular Conformation and Interactions. In P. A. Sandford & A. Laskin (Eds.), Extracellular Microbial Polysaccharides (Vol. 45, pp. 90–102). American Chemical Society. 10.1021/bk-1977-0045.ch007

Morris, E. R. (2019). Ordered conformation of xanthan in solutions and “weak gels”: Single helix, double helix – or both? Food Hydrocolloids, 86, 18–25. 10.1016/j.foodhyd.2017.11.036

Morris, E. R., Rees, D. A., Young, G., Walkinshaw, M. D., & Darke, A. (1977). Order-disorder transition for a bacterial polysaccharide in solution. A role for polysaccharide conformation in recognition between *Xanthomonas* pathogen and its plant host. Journal of Molecular Biology, 110(1), 1–16. 10.1016/S0022-2836(77)80095-8

Ndukwe, I. E., Black, I., Castro, C. A., Vlach, J., Heiss, C., Roper, C., & Azadi, P. (2024). Permethylation as a strategy for high-molecular-weight polysaccharide structure analysis by nuclear magnetic resonance—Case study of Xylella fastidiosa extracellular polysaccharide. Magnetic Resonance in Chemistry, 62(5), 370–377. 10.1002/mrc.5413

Niesen, F. H., Berglund, H., & Vedadi, M. (2007). The use of differential scanning fluorimetry to detect ligand interactions that promote protein stability. Nature Protocols, 2(9), 2212–2221. 10.1038/nprot.2007.321

Norton, I. T., Goodall, D. M., Frangou, S. A., Morris, E. R., & Rees, D. A. (1984). Mechanism and dynamics of conformational ordering in xanthan polysaccharide. Journal of Molecular Biology, 175(3), 371–394. 10.1016/0022-2836(84)90354-1

Ostrowski, M. P., La Rosa, S. L., Kunath, B. J., Robertson, A., Pereira, G., Hagen, L. H., Varghese, N. J., Qiu, L., Yao, T., Flint, G., Li, J., McDonald, S. P., Buttner, D., Pudlo, N. A., Schnizlein, M. K., Young, V. B., Brumer, H., Schmidt, T. M., Terrapon, N., … Martens, E. C. (2022). Mechanistic insights into consumption of the food additive xanthan gum by the human gut microbiota. Nature Microbiology, 7(4), 556–569. 10.1038/s41564-022-01093-0

Patel, J., Maji, B., Moorthy, N. S. H. N., & Maiti, S. (2020). Xanthan gum derivatives: Review of synthesis, properties and diverse applications. RSC Advances, 10(45), 27103–27136. 10.1039/D0RA04366D

Pelletier, E., Viebke, C., Meadows, J., & Williams, P. A. (2001). A rheological study of the order-disorder conformational transition of xanthan gum. Biopolymers, 59(5), 339–346. 10.1002/1097-0282(20011015)59:5%3C339::AID-BIP1031%3E3.0.CO;2-A

Petri, D. F. S. (2015). Xanthan gum: A versatile biopolymer for biomedical and technological applications. Journal of Applied Polymer Science, 132(23). 10.1002/app.42035

Rudolph, K. (1993). Infection of the plant by Xanthomonas. In J. G. Swings & E. L. Civerolo (Eds.), Xanthomonas (pp. 193–264). Springer Netherlands. 10.1007/978-94-011-1526-1_4

Salinas, S. R., Bianco, M. I., Barreras, M., & Ielpi, L. (2011). Expression, purification and biochemical characterization of GumI, a monotopic membrane GDP-mannose:glycolipid 4-β-d-mannosyltransferase from Xanthomonas campestris pv. Campestris. Glycobiology, 21(7), 903–913. 10.1093/glycob/cwr022

Sato, T., Kojima, S., Norisuye, T., & Fujita, H. (1984). Double-Stranded Helix of Xanthan in Dilute Solution: Further Evidence. Polymer Journal, 16(5), 423–429. 10.1295/polymj.16.423

Schäfer, A., Tauch, A., Jäger, W., Kalinowski, J., Thierbach, G., & Pühler, A. (1994). Small mobilizable multi-purpose cloning vectors derived from the Escherichia coli plasmids pK18 and pK19: Selection of defined deletions in the chromosome of Corynebacterium glutamicum. Gene, 145(1), 69–73. 10.1016/0378-1119(94)90324-7

Schmid, J., Huptas, C., & Wenning, M. (2016). Draft Genome Sequence of the Xanthan Producer Xanthomonas campestris LMG 8031. Genome Announcements, 4(5), e01069–16. 10.1128/genomeA.01069-16

Schmid, J., Sieber, V., & Rehm, B. (2015). Bacterial exopolysaccharides: Biosynthesis pathways and engineering strategies. Frontiers in Microbiology, 6, 496. 10.3389/fmicb.2015.00496

Shashkov, A. S., Lipkind, G. M., Knirel, Y. A., & Kochetkov, N. K. (1988). Stereochemical factors determining the effects of glycosylation on the^13^ C chemical shifts in carbohydrates. Magnetic Resonance in Chemistry, 26(9), 735–747. 10.1002/mrc.1260260904

Shatwell, K. P., Sutherland, I. W., Dea, I. C. M., & Ross-Murphy, S. B. (1990). The influence of acetyl and pyruvate substituents on the helix-coil transition behaviour of xanthan. Carbohydrate Research, 206(1), 87–103. 10.1016/0008-6215(90)84009-J

Smith, I. H., Symes, K. C., Lawson, C. J., & Morris, E. R. (1981). Influence of the pyruvate content of xanthan on macromolecular association in solution. International Journal of Biological Macromolecules, 3(2), 129–134. 10.1016/0141-8130(81)90078-7

Söderman, P., Jansson, P.-E., & Widmalm, G. (1998). Synthesis, NMR spectroscopy and conformational studies of the four anomeric methyl glycosides of the trisaccharide D-Glcp-(1→3)-[D-Glcp-(1→4)]-α-D-Glcp. Journal of the Chemical Society, Perkin Transactions 2, (3), 639–648. 10.1039/a707346a

Speciale, I., Notaro, A., Garcia-Vello, P., Di Lorenzo, F., Armiento, S., Molinaro, A., Marchetti, R., Silipo, A., & De Castro, C. (2022). Liquid-state NMR spectroscopy for complex carbohydrate structural analysis: A hitchhiker’s guide. Carbohydrate Polymers, 277, 118885. 10.1016/j.carbpol.2021.118885

Sutherland, I. W. (1998). Novel and established applications of microbial polysaccharides. Trends in Biotechnology, 16(1), 41–46. 10.1016/S0167-7799(97)01139-6

Tako, M., & Nakamura, S. (1984). Rheological Properties of Deacetylated Xanthan in Aqueous Media. Agricultural and Biological Chemistry, 48(12), 2987–2993. 10.1271/bbb1961.48.2987

Tomofuji, Y., Matsuo, K., & Terao, K. (2022). Kinetics of denaturation and renaturation processes of double-stranded helical polysaccharide, xanthan in aqueous sodium chloride. Carbohydrate Polymers, 275, 118681. 10.1016/j.carbpol.2021.118681

Torzewska, A., Kondakova, A. N., Perepelov, A. V., Senchenkova, S. N., Shashkov, A. S., Rozalski, A., & Knirel, Y. A. (2001). Structure of the O-specific polysaccharide of Proteus vulgaris O37 and close serological relatedness of the lipopolysaccharides of P. vulgaris O37 and P. vulgaris O46. FEMS Immunology & Medical Microbiology, 31(3), 227–234. 10.1111/j.1574-695X.2001.tb00524.x

Vorhölter, F.-J., Schneiker, S., Goesmann, A., Krause, L., Bekel, T., Kaiser, O., Linke, B., Patschkowski, T., Rückert, C., Schmid, J., Sidhu, V. K., Sieber, V., Tauch, A., Watt, S. A., Weisshaar, B., Becker, A., Niehaus, K., & Pühler, A. (2008). The genome of Xanthomonas campestris pv. Campestris B100 and its use for the reconstruction of metabolic pathways involved in xanthan biosynthesis. *Journal of Biotechnology*, *134*(1–2), 33–45. 10.1016/j.jbiotec.2007.12.013

Wang, X., Zheng, D., & Liang, R. (2016). An Efficient Electro-Competent Cells Generation Method of Xanthomonas campestris pv. campestris: Its Application for Plasmid Transformation and Gene Replacement. Advances in Microbiology, 6(2), 79–87. 10.4236/aim.2016.62008

Whitfield, C. (1988). Bacterial extracellular polysaccharides. Canadian Journal of Microbiology, 34(4), 415–420. 10.1139/m88-073

Whitfield, C. (2006). Biosynthesis and Assembly of Capsular Polysaccharides in *Escherichia coli*. Annual Review of Biochemistry, 75(1), 39–68. 10.1146/annurev.biochem.75.103004.142545

Wilson, D. B., Sahm, H., Stahmann, K.-P., & Koffas, M. (Eds.). (2020). In Industrial microbiology (pp. 279–281). Wiley-VCH.

Wu, M., Shi, Z., Ming, Y., Wang, C., Qiu, X., Li, G., & Ma, T. (2021). Thermostable and rheological properties of natural and genetically engineered xanthan gums in different solutions at high temperature. International Journal of Biological Macromolecules, 182, 1208–1217. 10.1016/j.ijbiomac.2021.05.008

Yao, H.-Y.-Y., Wang, J.-Q., Yin, J.-Y., Nie, S.-P., & Xie, M.-Y. (2021). A review of NMR analysis in polysaccharide structure and conformation: Progress, challenge and perspective. Food Research International, 143, 110290. 10.1016/j.foodres.2021.110290

