## Supplementary material for "NMR assignment and functional characterization of biotechnologically modified xanthan"

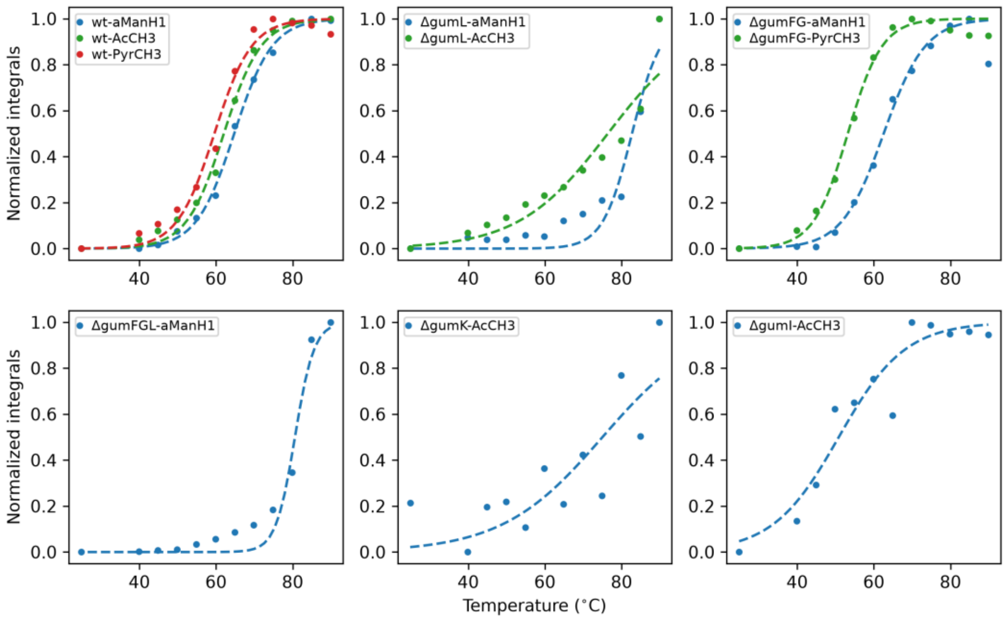


**Figure S1**: Fitting of the integrals of reporter signals to a sigmoidal function for $T_m$ determination. The results of fits are shown in Table 3.

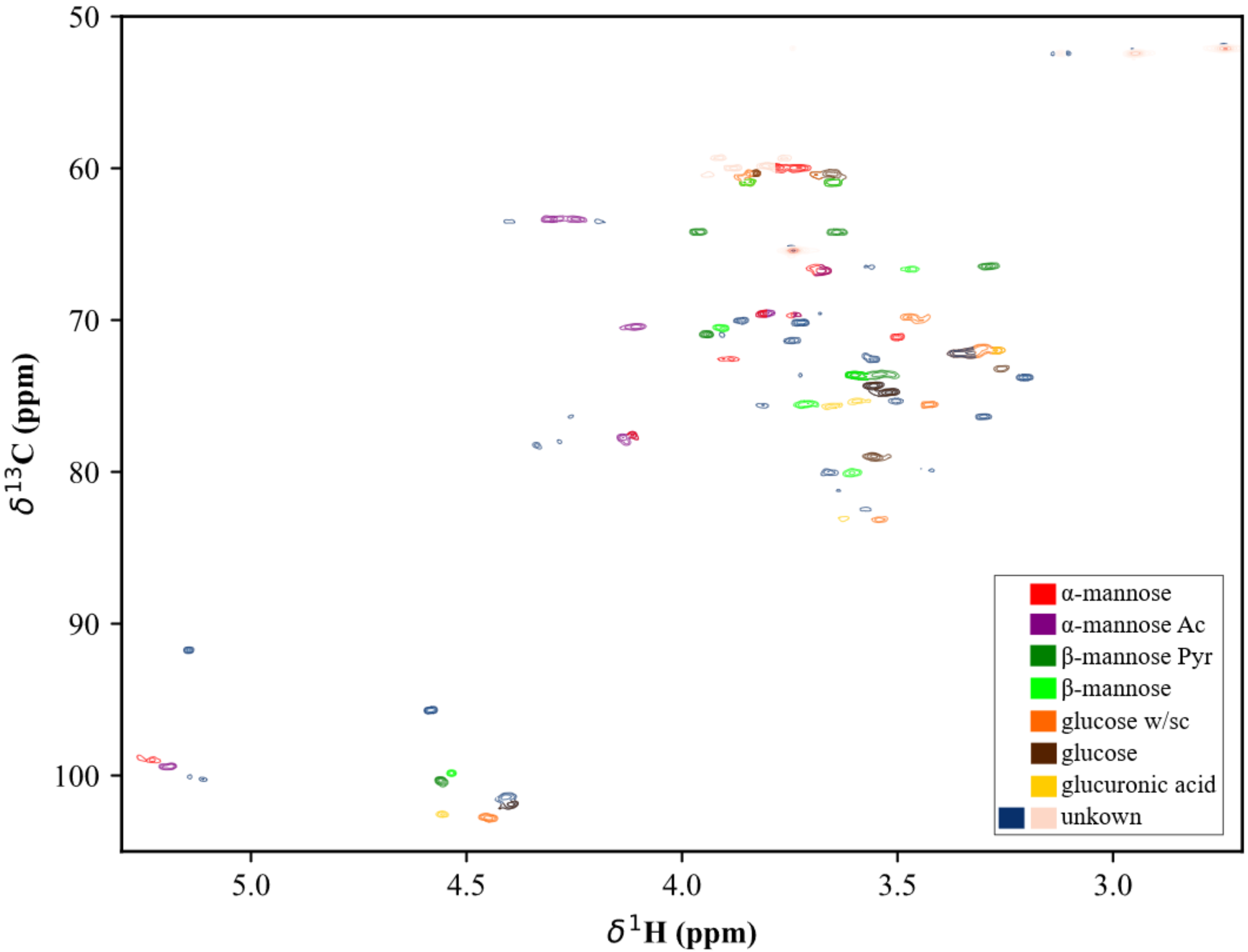


**Figure S2**: Magnified $^{1}$H $^{13}$C NMR chemical shift assignment of xanthan gum with assigned cross-peaks corresponding to the different sugar residues and their respective carbon–proton pairs. The signal assignments were based on published xanthan NMR data and additional spectra of the xanthan gum wt.

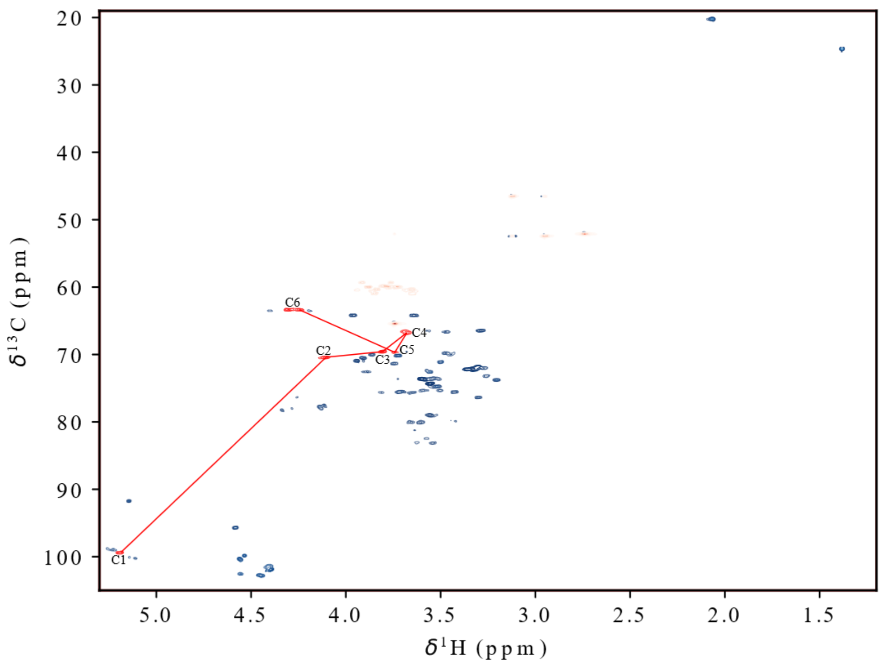


**Figure S3**: wt HSQC showing the individual spin system of the acetylated α-mannose, displaying the C/H resonances for each position from C1/H1 to C6/H6 and the correlations between them in red.

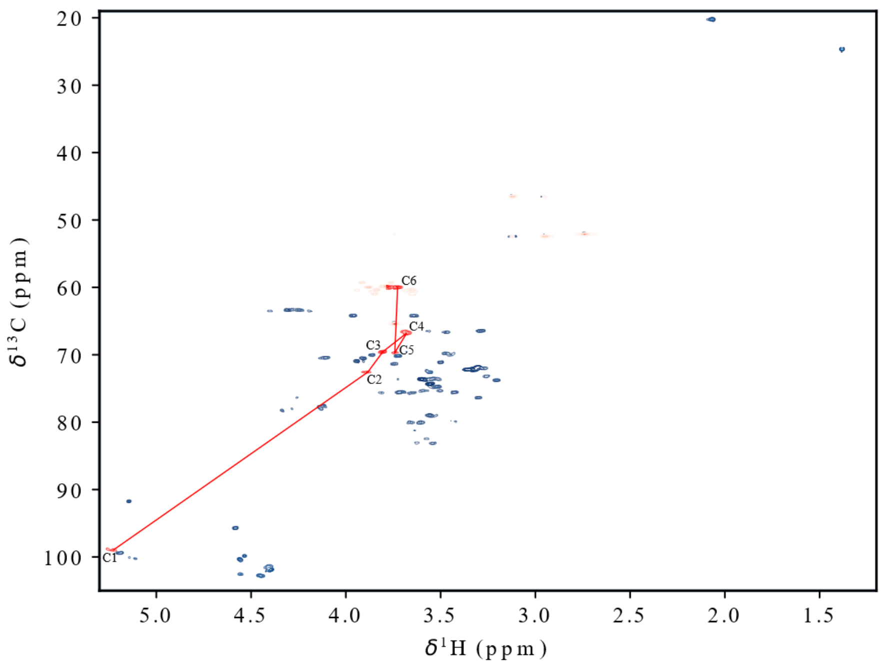


**Figure S4**: wt HSQC showing the individual spin system of the non-acetylated α-mannose, displaying the C/H resonances for each position from C1/H1 to C6/H6 and the correlations between them in red.

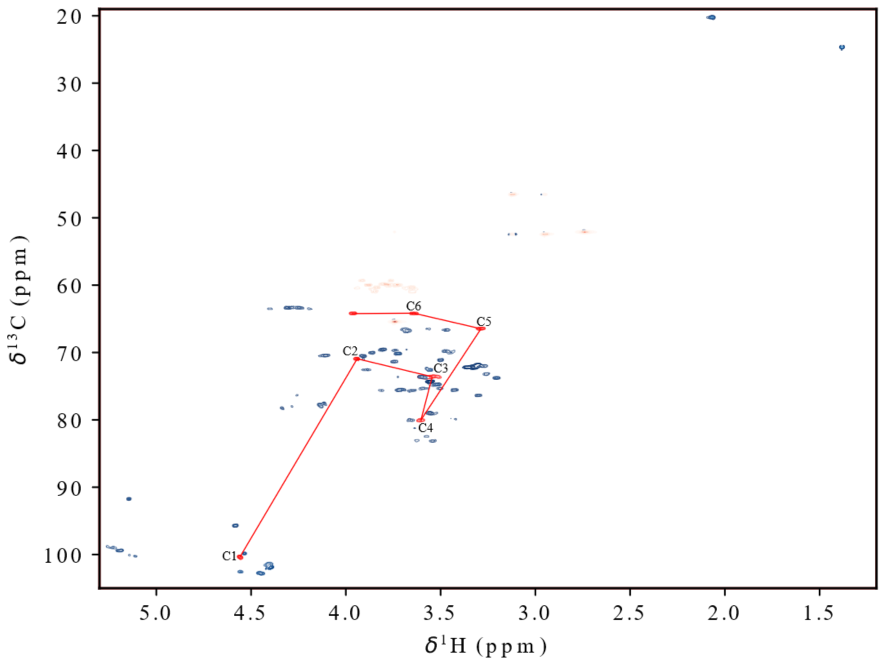


**Figure S5**: wt HSQC showing the individual spin system of the pyruvylated β-mannose, displaying the C/H resonances for each position from C1/H1 to C6/H6 and the correlations between them in red.

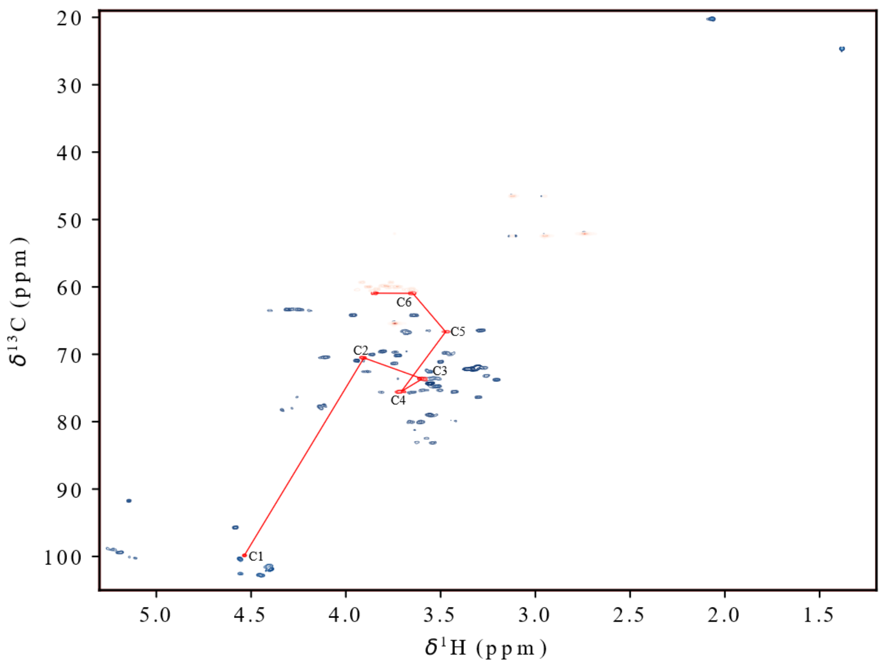


**Figure S6**: wt HSQC showing the individual spin system of the non-pyruvylated β-mannose, displaying the C/H resonances for each position from C1/H1 to C6/H6 and the correlations between them in red.

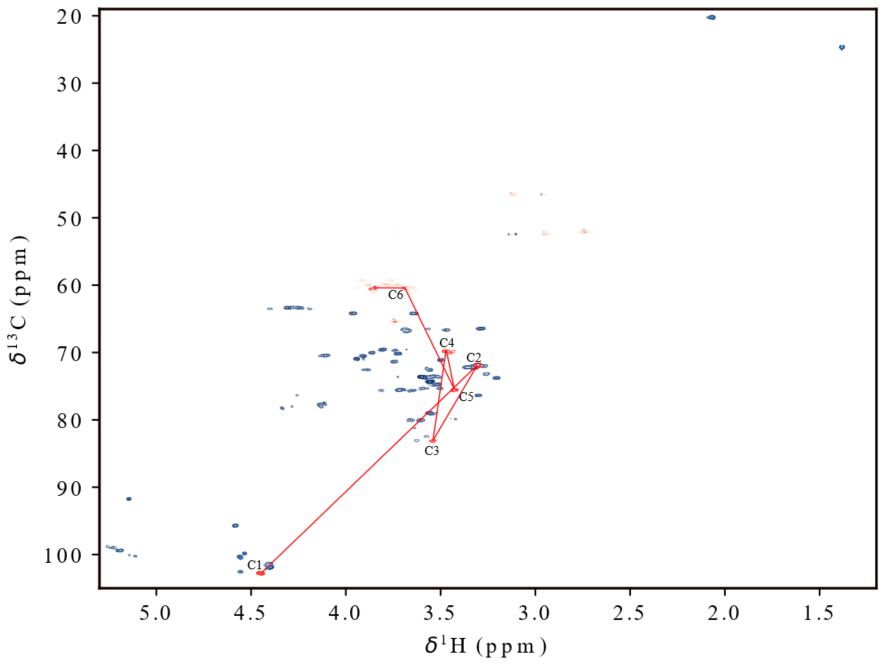


**Figure S7**: wt HSQC showing the individual spin system of the side-chain-linked glucose, displaying the C/H resonances for each position from C1/H1 to C6/H6 and the correlations between them in red.

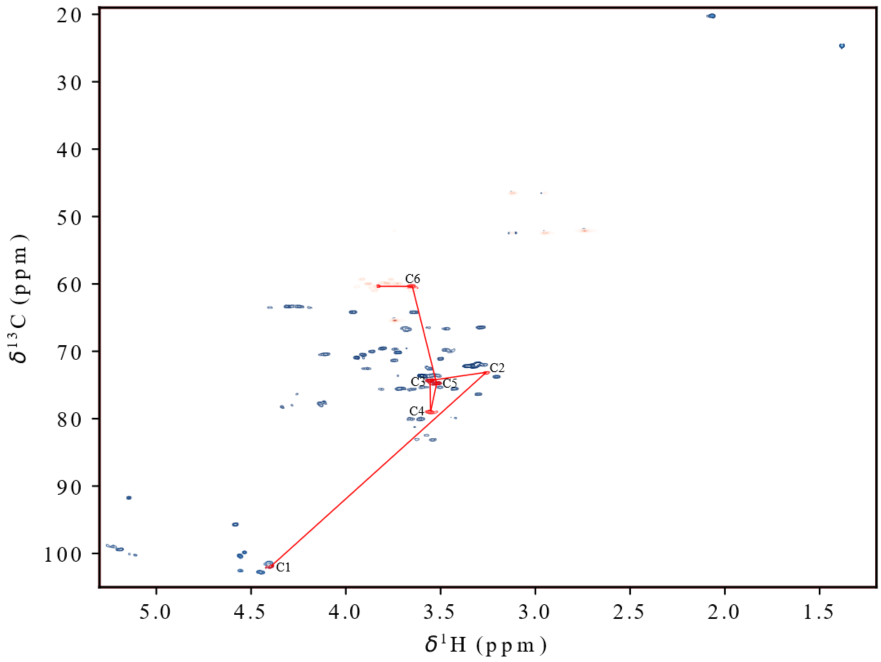


**Figure S8**: wt HSQC showing the individual spin system of the glucose without side chain, displaying the C/H resonances for each position from C1/H1 to C6/H6 and the correlations between them in red.

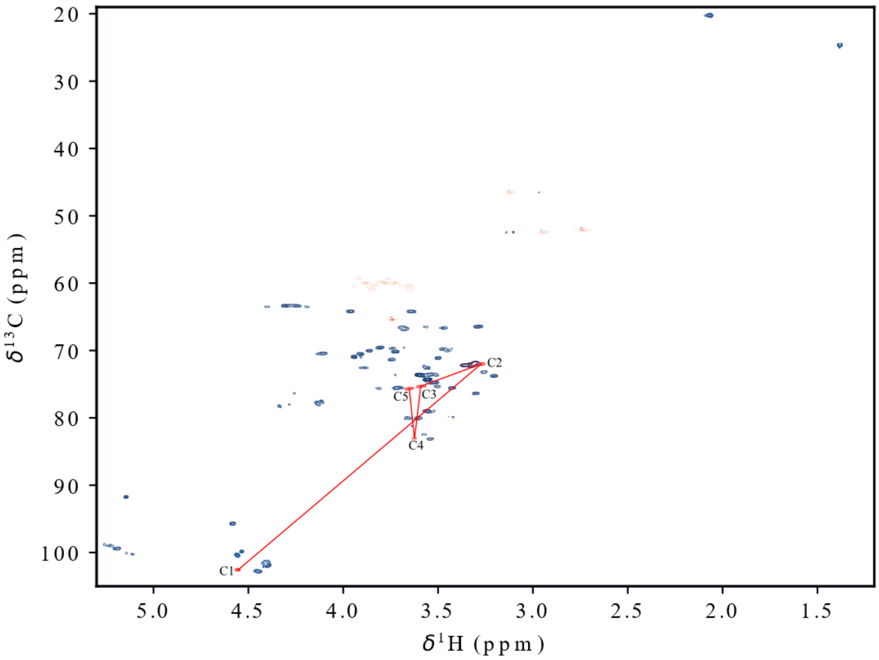


**Figure S9**: wt HSQC showing the individual spin system (without C6) of the glucuronic acid, displaying the C/H resonances for each position from C1/H1 to C5/H5 and the correlations between them in red.

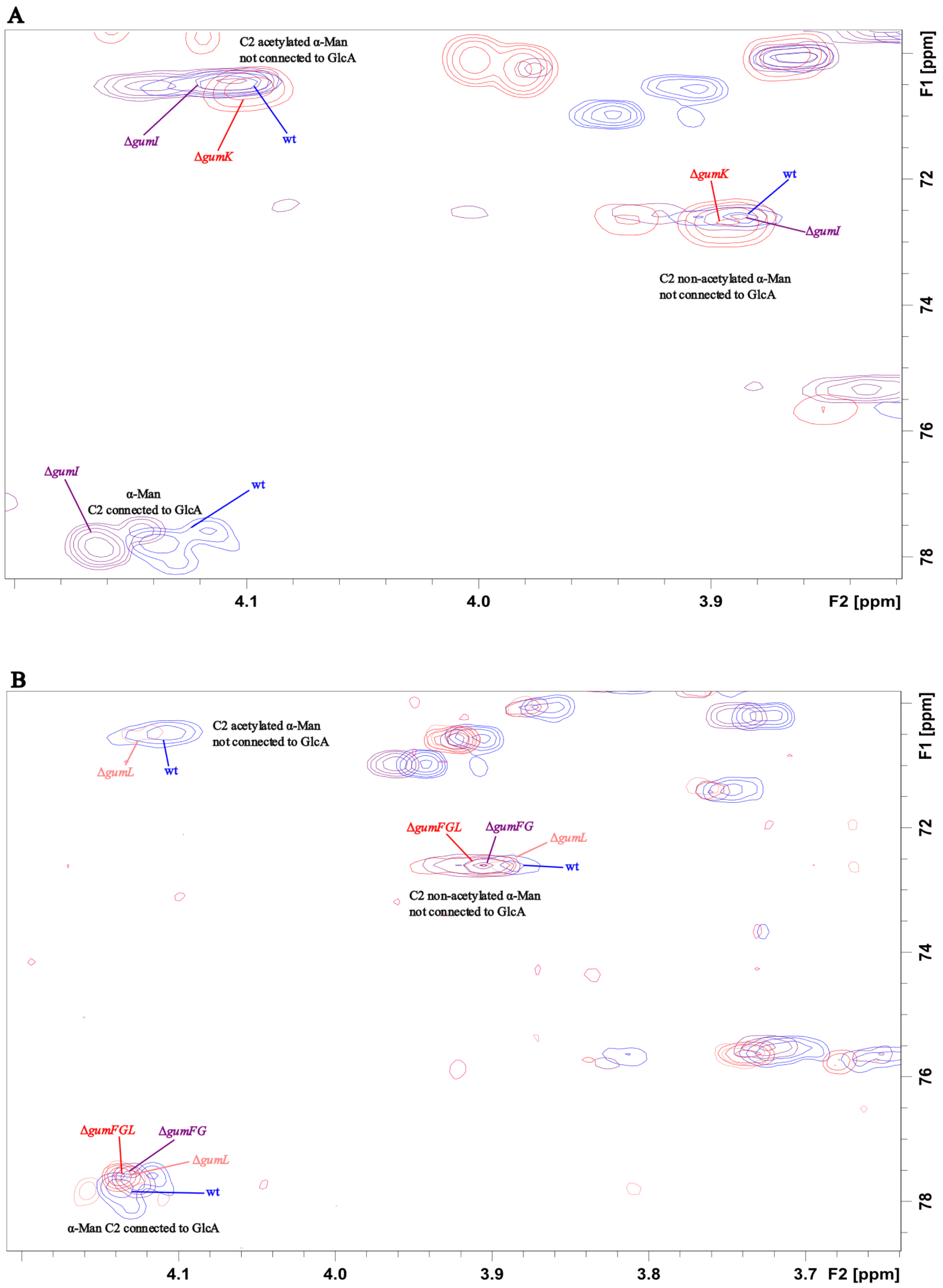


**Figure S10**: Overlay of the HSQC spectra of wild-type and xanthan variants in the region of the α-Man C2 resonance. (A) Overlay of the wild-type spectrum with the spectra of Δ*gumI* and Δ*gumK*. (B) Overlay of the wild-type spectrum with the spectra of Δ*gumFG*, Δ*gumL*, and Δ*gumFGL*.

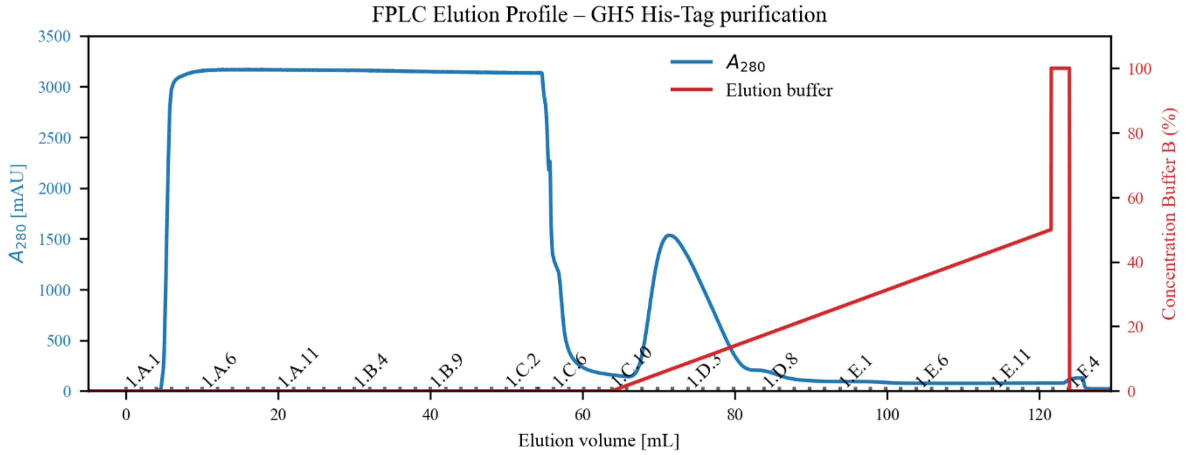


**Figure S11**: Elution profile of the RuGH5a His-tag purification. Fractions were collected into a 96-deep-well plate; the lower part of the x-axis indicates the elution volume, the upper part of the x-axis indicates the well position, and the y-axis the absorbance at 280 nm (mAU). The red line indicates the concentration of buffer B used for elution.

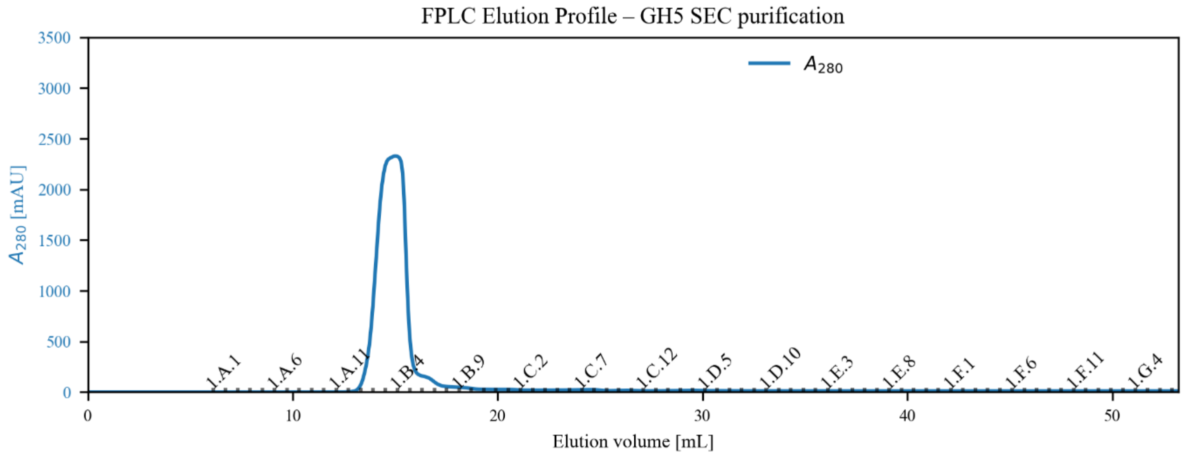


**Figure S12**: Elution profile of the RuGH5a size-exclusion purification. Fractions were collected into a 96-deep-well plate; the lower part of the x-axis indicates the elution volume, the upper part of the x-axis indicates the well position, and the y-axis the absorbance at 280 nm (mAU).

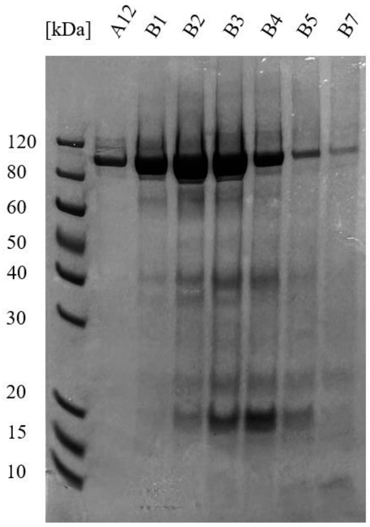


**Figure S13**: 12% SDS-Page of RuGH5a after SEC purification.

### Plasmids

pK19GmΔ*gumK*
pK19GmΔ*gumI*
pK19GmΔ*gumF*
pK19GmΔ*gumG*
pK19GmΔ*gumL*

### Primers

Table S1: All primers used to create the strains capable of producing the xanthan variants.

| **Oligo** | **5'→3'** |
|---|---|
| *gumFG*_check_fw | CATGATTGTGGCATCCGACG |
| *gumFG*_check_rev | TGCCCTGCATACTTCTCCAC |
| *gumFG*_fld_fw_OE | ATACGGTGACAGGGGCATCGCAGAATCATCAG-TCGATGTG |
| *gumFG*_fld_rev_BamHI | TATATAGGATCCCGAACAGATCGCCGTCATTC |
| *gumFG*_flu_fw_XbaI | TATATATCTAGAGCCGTTATTGAAACGGATGG |
| *gumFG*_flu_rev_OE | CACATCGACTGATGATTCTGCGATGCCCCTGTCAC-CGTAT |
| *gumL*_check_fw | ACGTCTTCCATGTAGGTCAC |
| *gumL*_check_rev | CGGAGAGAAAATCCAGCAAG |
| *gumL*_fld_fw_OE | GTTTGAAGGAGGATCCCTGTAAC-GACAATGCATGGCCAGC |
| *gumL*_fld_rev_HindIII | TATATAAAGCTTGTTGCCGAAGGCCACCAAC |
| *gumL*_flu_fw_XbaI | TATATATCTAGAGTATGCCGAAGGCATCCATG |
| *gumL*_flu_rev_OE | GCTGGCCATGCATT-GTCGTTACAGGGATCCTCCTTCAAAC |

| | |
|---|---|
| *gumF*_fld_fw_OE | ATACGGTGACAGGGGCATCGATGACGAC-GGCTGCGATC |
| *gumF*_fld_rev_NheI | TATATAGCTAGCCAACCAGCAAACTGCAGACC |
| *gumF*_flu_rev_OE | GATCGCAGCCGTCGTCATCGATGCCCCTGTCACCG-TAT |
| *gumG*_fld_fw_OE | CCTCAACTGTCGCGGAGCGCCAGAATCATCAG-TCGATGTG |
| *gumG*_flu_fw_XbaI | TATATATCTAGAATTGTTCTGGGGCCTGGATG |
| *gumG*_flu_rev_OE | CACATCGACTGATGATTCTGGCGCTCCGCGACAGTT-GAG |
| *gumF*_gDNA_check_rev | GCAGCACATCCAGTGCAAAC |
| *gumG*_gDNA_check_fw | GGCTGTCGCAGTTGTTCTC |
| *gumF*_GA_fw | CACACAGGAAACAGCAGGTGAATAC-GGTGACAGGGGC |
| *gumF*_GA_rev | TCGAATTTGCTTTCGAATTGTCATGCCGACACCG-GACGTG |
| pSRK_check_fw | TGCTTCCGGCTCGTATGTTG |
| pSRK_check_rev | TTAACGACCCTGCCCTGAAC |
| pSRK_BamHI_fw | TATATATGGATCCAATTCGAAAGCAAATTCGAA |
| pSRK_BamHI_rev | TATATAGGATCCTGCTGTTTCCTGTGTGAAA |
| *gumK*_KOproof_fw | ATCGCCTGCATTTTTGTTGGC |
| *gumK*_KOproof_rev | AACTGCTGATGACAAATTCCGC |
| GA_gumK_1 | ATGAGCGTCTCTCCCGCAGCTC |
| GA_gumK_2 | GAGCTGCGGGAGAGACGCTC |
| *gumK*_fld_fw_OE | GGAGATATGCGAGGAGGCGCTTTTTTGATC |
| *gumK*_fld_rev_HindIII | TATATAAAGCTTTGTACTTCTCAACCGGCTCG |
| *gumK*_flu_fw_XbaI | TATATATCTAGACTGGTCTGGATCGGCAAGATC |
| *gumK*_flu_rev_OE | GCCTCCTCGCATATCTCCCTTCATGCGG |
| *gumI*_KO_proof_fw | AAACTCATCGGTACACCTGTCTGGG |
| *gumI*_KO_proof_rev | TGATCTTGCCGATCCAGACCAGC |
| *gumI*_fld_fw_OE | GCAGGAGGCCGTGCGATGACCGCAGAGACATCGAC-CATG |
| *gumI*_fld_rev_HindIII | TATATAAAGCTTCATGCGCCAGCGATAATCAG |
| *gumI*_flu_fw_XbaI | TATATATCTAGAGCTTGAATTGCTGCAGGCTG |
| *gumI*_flu_rev_OE | CATGGTCGATGTCTCTGCGGTCATCGCAC-GGCCTCCTGC |
